# Loss of noradrenergic *Fkbp5* disrupts social behavior and norepinephrine dynamics in the basolateral amygdala

**DOI:** 10.64898/2025.12.10.693477

**Authors:** Joeri Bordes, Leona Stont, Thomas Bajaj, Simon Chang, Tim Ebert, Lucas Miranda, Patrick Schlegel, Maya Reinhardt, Max Pöhlmann, Yara Mecdad, Lotte van Doeselaar, Huanqing Yang, Veronika Kovarova, Shiladitya Mitra, Meri De Angelis, Bertram Müller-Myhsok, Jan M. Deussing, Anna Beyeler, Nils C. Gassen, Mathias V. Schmidt

## Abstract

Social dysfunction is common in depression and varies with stress exposure and genetic risk. The current study identifies a cell-type specific role for *Fkbp5*, a glucocorticoid receptor co-chaperone, in noradrenergic neurons engaged during social stress. Acute social stress upregulated *Fkbp5* in the locus coeruleus (LC), whereas repeated exposure attenuated this effect. Noradrenergic *Fkbp5* deletion (*Fkbp5*^Nat^) increased pro-social behavior exclusively in male mice. In the basolateral amygdala (BLA), social interaction reduced norepinephrine (NE) turnover in wild-type but not *Fkbp5*^Nat^ mice. Consistently, proteomics revealed mitochondrial/energy and synapse-related remodeling in BLA neurons. Miniscope imaging showed that behavior-locked NE transients in BLA were selectively blunted in *Fkbp5*^Nat^ mice during interaction with outbred CD1 conspecifics, while same-strain C57BL/6N encounters preserved NE dynamics. Together, this study indicates that *Fkbp5* tunes LC-BLA output to social salience in a sex- and context-dependent manner, suggesting a circuit-specific route to normalize social salience without broadly suppressing noradrenergic function.

## Introduction

Social interactions are fundamental to daily life and critical for a range of societal outcomes, from forming friendships to career success. Yet, the expression of social behavior and response to social challenges show high inter-individual variability, reflecting a complex interplay of emotional and cognitive processes^1^. Abnormal social behavior is a core characteristic of many psychiatric disorders, notably major depressive disorder (MDD)^2–5^. MDD is frequently associated with profound social dysfunction, which can manifest as social withdrawal^6,7^, but also maladaptive behaviors like aggression^8,9^. These social impairments are a recognized risk factor for disease persistence and symptom relapse^10,11^. Notably, social dysfunction in MDD has been identified as a semi-independent domain, often persisting even after the remission of other core depressive symptoms^11,12^.

The vulnerability to MDD is shaped by a complex interplay of genetics, epigenetics, and the environment^13,14^. Human genetic studies have implicated polymorphisms in the *FKBP5* gene in susceptibility to MDD ^15–17^, particularly in individuals exposed to severe stress^18–20^. FKBP51, encoded by the *FKBP5* gene, is an important regulator of the hypothalamic-pituitary-adrenal (HPA) axis and stabilizes the glucocorticoid receptor (GR)-complex structure, thereby decreasing the binding to glucocorticoids and hampering the nuclear translocation of the GR complex^14,21,22^. Beyond its role as a co-chaperone at the GR complex, *FKBP5*/FKBP51 contributes to protein homeostasis by engaging the autophagy machinery and priming autophagic flux, a process that has also been associated with antidepressant response in preclinical and clinical studies^23^. While the link between *FKBP5*, stress, and HPA axis dysregulation is well-established, its region-specific effects on social function remain largely unknown. Preclinical mouse models have shown that *Fkbp5* is highly expressed and regulated in key stress-related brain regions, including the hypothalamus, amygdala, hippocampus, bed nucleus of stria terminalis, dorsal raphe nucleus (DRN), and locus coeruleus (LC)^24–26^. Notably, the LC-noradrenergic system represents a prime candidate for a mechanism that links stress-related genetic risk to social and emotional dysfunction, since this region (1) shows high baseline *Fkbp5* expression^24^, (2) plays a crucial role in the stress response system by initiating the sympathetic–adrenal–medullary (SAM) axis, and (3) has connections with the HPA axis via CRF neurons from several stress-related brain regions that can further innervate the LC^27–29^.

The LC is strongly activated by stress exposure and has widespread noradrenergic projections throughout the brain, orchestrating large-scale network processes influencing many cognitive functions^30,31^. In addition, the LC has pathway-specific influences. Notably, LC-induced norepinephrine (NE) release in the basolateral amygdala (BLA) increases anxiety-like behavior^32^. Several studies have shown that the LC strongly responds to social stress exposure, via the general activation of the LC and adapting specific projection circuits^33,34^, however the direct influence of the LC on social behavior remains to be uncovered.

The social behavioral construct encompasses a wide spectrum of distinct behaviors^35^, making its investigation a considerable technological challenge^36–38^. Traditional behavioral tasks often rely on oversimplified constructs that fail to capture the full richness of social interactions^38^. Recent advancements in automated behavioral phenotyping have allowed for high-throughput analysis using pose estimation^39,40^ and subsequently automated supervised behavioral classification^41^. These tools provide a powerful mean to meticulously quantify the complex social behaviors influenced by *Fkbp5* and the noradrenergic system, enabling a deeper mechanistic understanding.

In the current study, we aimed to unravel the region-specific role of *Fkbp5* regulation on the neurobiological behavioral changes induced by stress exposure. We employed the open-source Python package DeepOpenField (DeepOF) to perform deep phenotyping of social behaviors and enhance the understanding of stress exposure impact on social behaviors^41,42^. We show that acute social challenge selectively modulates LC activity and induces *Fkbp5* expression. Noradrenergic Fkbp5 deletion produces a male-specific alteration in social behavior accompanied by persistent changes in BLA NE signaling and related molecular pathways. To test context, we compared interaction with a novel, outbred CD1 conspecific versus an unfamiliar same-strain C57BL/6N; the NE deficit emerged selectively during CD1 encounters, indicating strain-dependent social salience. Taken together, our findings identify a previously unrecognized, noradrenergic cell-type-specific role for *Fkbp5* in shaping social behavior and provide a compelling mechanistic link between genetic vulnerability, stress, and sex-specific social dysfunction.

## Results

### *Fkbp5* shows a social stress-specific upregulation in the LC

The expression patterns of *Fkbp5* were examined in the LC and DRN brain regions across various chronic and acute stress paradigms. Stress-naïve male mice were euthanized under baseline conditions, nonstressed (NS) or exposed to an acute stressor before euthanasia four hours after intervention, which was either acute social defeat stress (ASDS), or restraint stress (Res). Another group of animals underwent 21 days of chronic social defeat stress (CSDS) followed by either no additional stressor (CSDS, euthanasia 24 hours after CSDS finished), or following acute stress exposure, which was either acute social defeat stress (CSDS-ASDS), or restraint stress (CSDS-Res) four hours prior to euthanasia, as depicted in Figure 1A.

**Figure 1.**
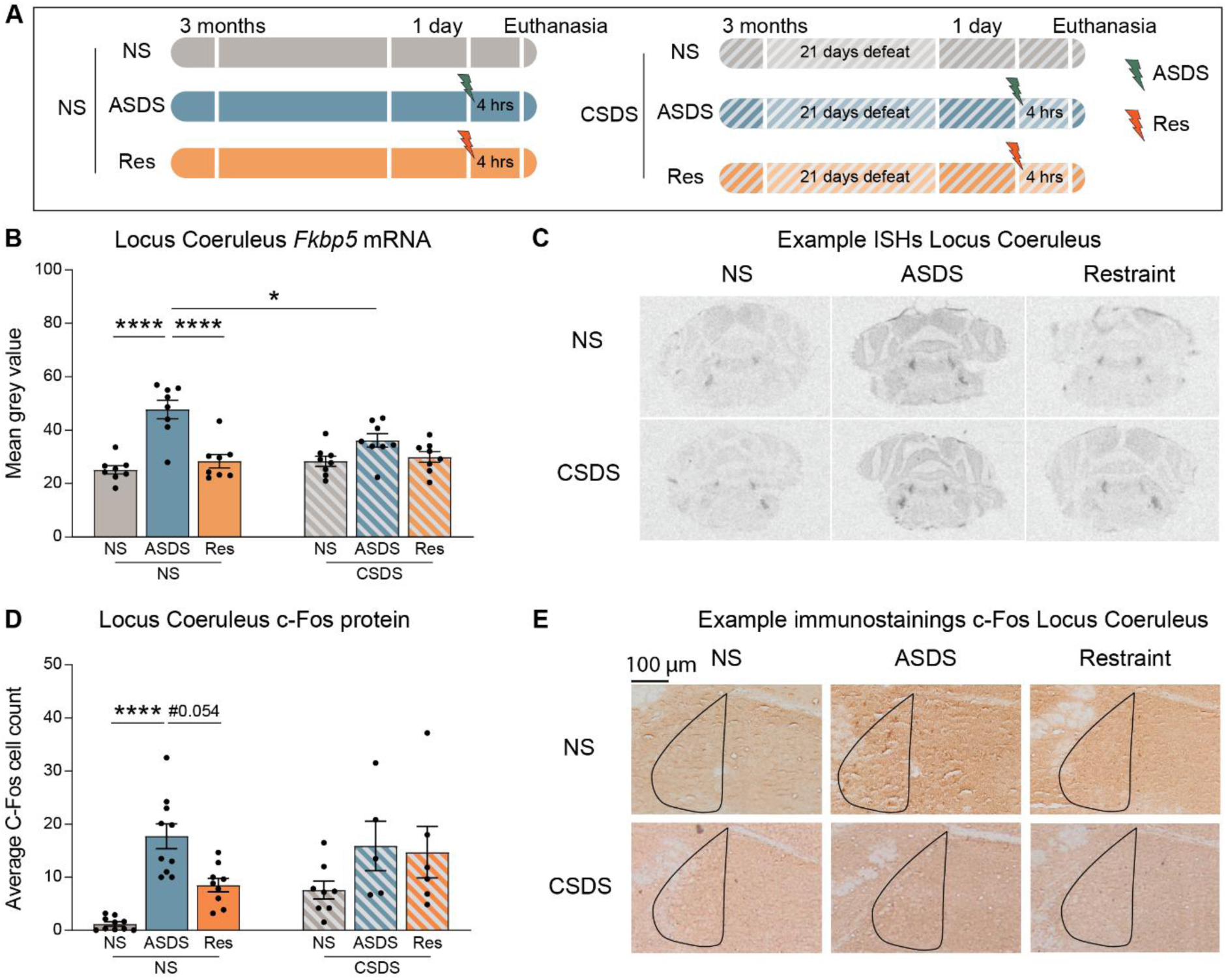
*Fkbp5* and c-Fos expression in the Locus Coeruleus. **A)** Overview of the different acute and chronic stress conditions. **B)** Fkbp5 mRNA expression in the LC revealed a significant interaction effect using a two-way ANOVA (Acute×Chronic) (F(2,42)=5.696, p=0.0065) and a main effect of Acute (F(2,42)=23.06, p=1.7e-7), with no main effect of Chronic (F(1,42)=1.325, p=0.256). Post-hoc Tukey revealed that *Fkbp5* mRNA expression in the LC was significantly increased in NS-ASDS compared to NS-NS (p=6.9^e-7^) and NS-Res (p=5.47^e-6^). The NS-ASDS also had significantly increased *Fkbp5* levels compared to CSDS-ASDS (p=0.017). The CSDS-ASDS mice did have altered *Fkbp5* levels compared to CSDS-NS (p=0.22), or CSDS-Res (p=0.99). **C)** Example in-situ hybridization scans in the LC. **D)** c-Fos protein expression in the LC showed a significant main effect of “Acute” using a two-way ANOVA (Acute×Chronic) (F(2,42)=13.11, p=3.8e-5), with no Acute×Chronic interaction (F(2,42)=1.685, p=0.198) and no main effect of Chronic (F(1,42)=3.127, p=0.084). Post-hoc Tukey indicated higher c-Fos in NS-ASDS vs NS-NS (p=3.6e−5) and a trend toward higher levels vs NS-Res (p=0.054). NS-ASDS did not differ from CSDS-ASDS (p=0.996). Within CSDS, ASDS did not differ from NS (p=0.284) or Res (p=0.999). **E)** Example immunostaining in the LC. In panel B; n=8 for all groups. In panel D; n=10 for NS-NS and NS-ASDS, n=9 for NS-Res, n=8 for CSDS, n=5 for CSDS-ASDS, n=6 for CSDS-Res.

In mice without a history of chronic stress, acute social defeat stress led to a significant increase in *Fkbp5* mRNA levels compared to stress-naïve and acutely restrained mice in both the LC (Figure 1B-C) and DRN (Supplemental Figure 1A-B) regions. A similar effect was observed in mice with a history of chronic stress in the DRN region, where mice exposed to CSDS plus acute social defeat stress exhibited elevated *Fkbp5* mRNA levels compared to those in CSDS baseline conditions and CSDS plus acutely restrained conditions (Supplemental Figure 1A-B). However, the LC region displayed a different response after chronic stress, as mice exposed to CSDS plus acute social defeat stress did not show elevated *Fkbp5* mRNA levels compared to those in CSDS baseline conditions and CSDS plus acutely restrained conditions. Furthermore, the baseline acute social defeat stress mice demonstrated significantly higher *Fkbp5* mRNA levels compared to animals with a chronic stress history plus acute social defeat stress mice (Figure 1B-C).

The expression levels of the c-Fos protein in the LC exhibited a similar pattern to *Fkbp5* expression under acute stress conditions. Exposure to acute social defeat stress significantly upregulated c-Fos expression compared to stress-naïve, and a trend to acutely restrained mice (Figure 1D-E). In contrast to the *Fkbp5* results, a history of CSDS exposure did not result in altered c-Fos expression pattern between mice exposed to CSDS plus acute social defeat stress and those in CSDS baseline conditions or CSDS plus acutely restrained conditions (Figure 1D-E). Additionally, the baseline acute social defeat stress mice did not exhibit altered c-Fos expression compared to animals with a chronic stress history plus acute social defeat stress (Figure 1D-E).

### Conditional knockout of *Fkbp5* in the LC-NE system alters the social behavioral profile in male mice

Given the high expression of Fkbp5 in the LC and its selective induction by acute social stress, we next investigated its role during a social stress task with lower intensity: social interaction (SI) with an unfamiliar, unaggressive juvenile outbred CD1 conspecific in a novel environment. We generated a conditional *Fkbp5* knockout (KO) line using a driver line expressing Cre recombinase controlled by the noradrenergic-specific Nat promoter. This conditional knockout line was employed to investigate the precise effects of *Fkbp5* knockout on the noradrenergic system, including the LC (Figure 2A). To confirm the efficacy of the knockout, an RNAscope *in situ* hybridization study was performed using a co-expression analysis of *Fkbp5* and tyrosine hydroxylase (TH), a marker of noradrenergic cells. Quantitative analysis revealed a significant reduction in the percentage of *Fkbp*5-positive TH+ cells in the LC of *Fkbp5*^Nat^ (KO) mice compared to *Fkbp*5^lox/lox^ (WT) mice (Figure 2B–C).

**Figure 2.**
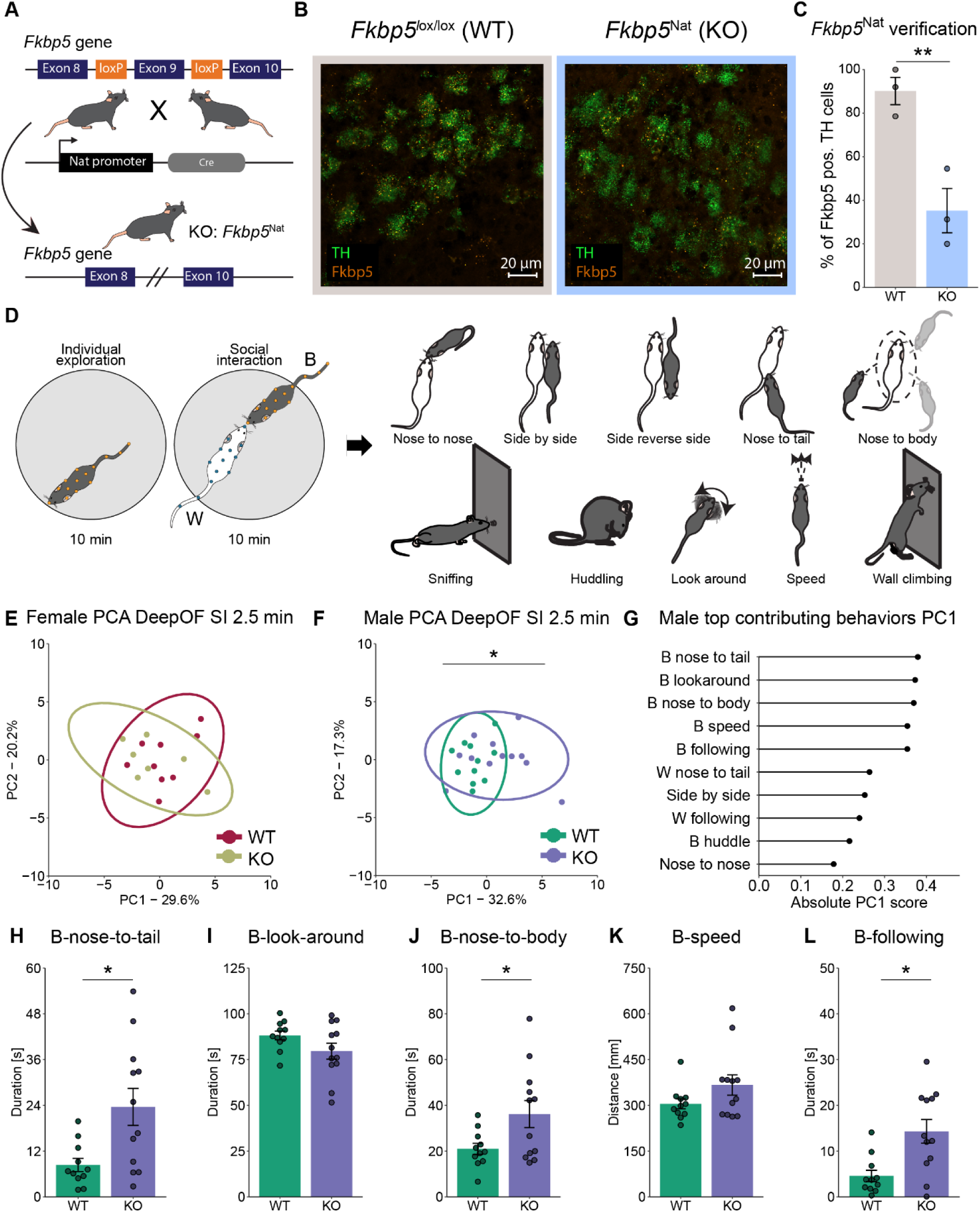
*Fkbp5*^Nat^ males show an altered social behavioral profile. **A)** Schematic overview of the generation of *Fkbp5* knockout in the noradrenergic cells. **B)** Representative RNAscope confocal images of *Fkbp5* mRNA expression in TH neurons within the LC. C) RNAscope *Fkbp5* quantification in the TH neurons revealed a significant reduction of the percentage of *Fkbp5* positive TH cells in KO animals (Two-tailed independent samples-t-test: T(4)=4.6, p=0.01). **D)** Schematic overview of the social interaction task and DeepOF identified individual and social behaviors. **E)** The female SI 2.5 min time bin PCA showed no difference in the PC1 eigenvalues between conditions. The PCA data consisted of all the SI DeepOF behavioral classifiers. Two-tailed independent samples t-test: T(14)=-0.72, p=0.49. **F)** The male SI 2.5 min time bin PCA showed a significant difference in the PC1 eigenvalues between conditions. Welch test: We(14.96)=-2.32, p=0.035. **G)** The top contributing behaviors for males in the SI 2.5 min time bin PC1 using the corresponding rotated loading scores. The top five behaviors were listed as potentially relevant behaviors for identifying genotype effect (B-nose-to-tail (0.38), B-look-around (–0.37), B-nose-to-body (0.37), B-speed (0.35), B-following (0.35). “B-” indicates C57Bl/6N behaviors and “W-” indicates CD1 behaviors. **H)** The 2.5 min duration of B-nose-to-tail. Wilcoxon test: Wx=27, p=0.016. **I)** The 2.5 min duration of B-look-around. Welch test: We(16,90)=1.71, p=0.11. **J)** The 2.5 min duration of B-nose-to-body. Wilxocon test: Wx=38, p=0.091. **K)** The 2.5 min duration of B-speed. Wilcoxon test: Wx=47, p=0.26. **L)** The 2.5 min duration of B-following. Wilcoxon test: Wx=25, p=0.011. In panel C; n=3 for WT and n=3 for KO. In panel D; n=9 for WT and n=7 for KO. In panels E-K; n=11 for WT and n=12 for KO. Sample size: panels B-C: WT: n=3, and KO: n=3. Panel E: WT: n=9, KO: n=7. Panels F-L: WT: n=11, and KO: n=12.

The social behavioral profile was examined during the SI task in both male and female mice (Figure 2D), utilizing four distinct time bins of 2.5 minutes each. A Principal Component Analysis (PCA) was employed to explore the social behavior profile across different time bins, including both sexes and genotypes. The PCA results indicated a disparity between the first 2.5-minute bin and all subsequent time bins, suggesting that the initial 2.5 minutes were most crucial for social behavioral phenotyping (Supplemental Figure 2A-B), in line with our previous findings^41^. To assess the social behavioral phenotype across sex, a PCA was conducted using the first 2.5-minute SI bin between female and male DeepOF social behaviors, regardless of genotype. A significant separation in the PCA was observed between female and male mice (Supplemental Figure 2C-D), indicating a different social behavioral profile across sexes. Therefore, further analyses were performed separately for each sex.

Separate PCA’s were conducted for the first 2.5-minute bin in female and male SI data to compare genotypes. In the female PCA, no significant differences were found between genotypes (Figure 2E). However, in the male PCA, a significant difference was observed (Figure 2F). Further examination of the male PCA data was performed via the exploration of the top contributing behaviors in PC1, determined by the corresponding rotated loading scores (Figure 2G). The top five contributing behaviors displayed potentially relevant patterns for identifying genotype effects, while other behaviors within the top 10 either contributed to the CD1 animal (“W-” behaviors) or had low rotated loading scores. Further analysis of the top contributing behaviors in the male 2.5-minute SI data revealed a significant increase in KO mice compared to WT mice in the expression of B-nose-to-tail, B-nose-to-body, and B-following behaviors, whereas no genotype difference was observed in B-look-around and B-speed behaviors (Figure 2H-L). To investigate the specificity of acute effects of social exposure, male mice were examined for their social behavioral profile following CSDS exposure. The PCA results did not indicate any alterations between genotypes, suggesting a specificity of the FKBP5-NE system to novel social interactions rather than repeated chronic social stress exposure (Supplemental Figure 2E-F).

### Social interaction drives BLA proteomic remodeling in *Fkbp5*^Nat^ mice

Given that *Fkbp5* deletion was restricted to noradrenergic neurons, we hypothesized that the noradrenergic projections might be altered in KO mice following SI. To test this, total NE content was quantified in microdissected tissue from major LC projection target areas, namely the BLA, the medial prefrontal cortex (mPFC), dorsal hippocampus (dHipp) and ventral hippocampus (vHipp), under NI and SI conditions. In the BLA, a significant increase in NE concentration was observed in WT_SI compared to WT_NI, an effect that was absent in KO mice (Figure 3H). These results suggest a differential functional activation of the LC-BLA circuit during social engagement between WT and KO mice. In the other assessed regions, no significant genotype- or interaction-dependent differences in total NE content were detected, showing a high specificity of LC Fkbp5 modulating NE output following SI in the BLA (Supplemental Figure 4A-D).

**Figure 3.**
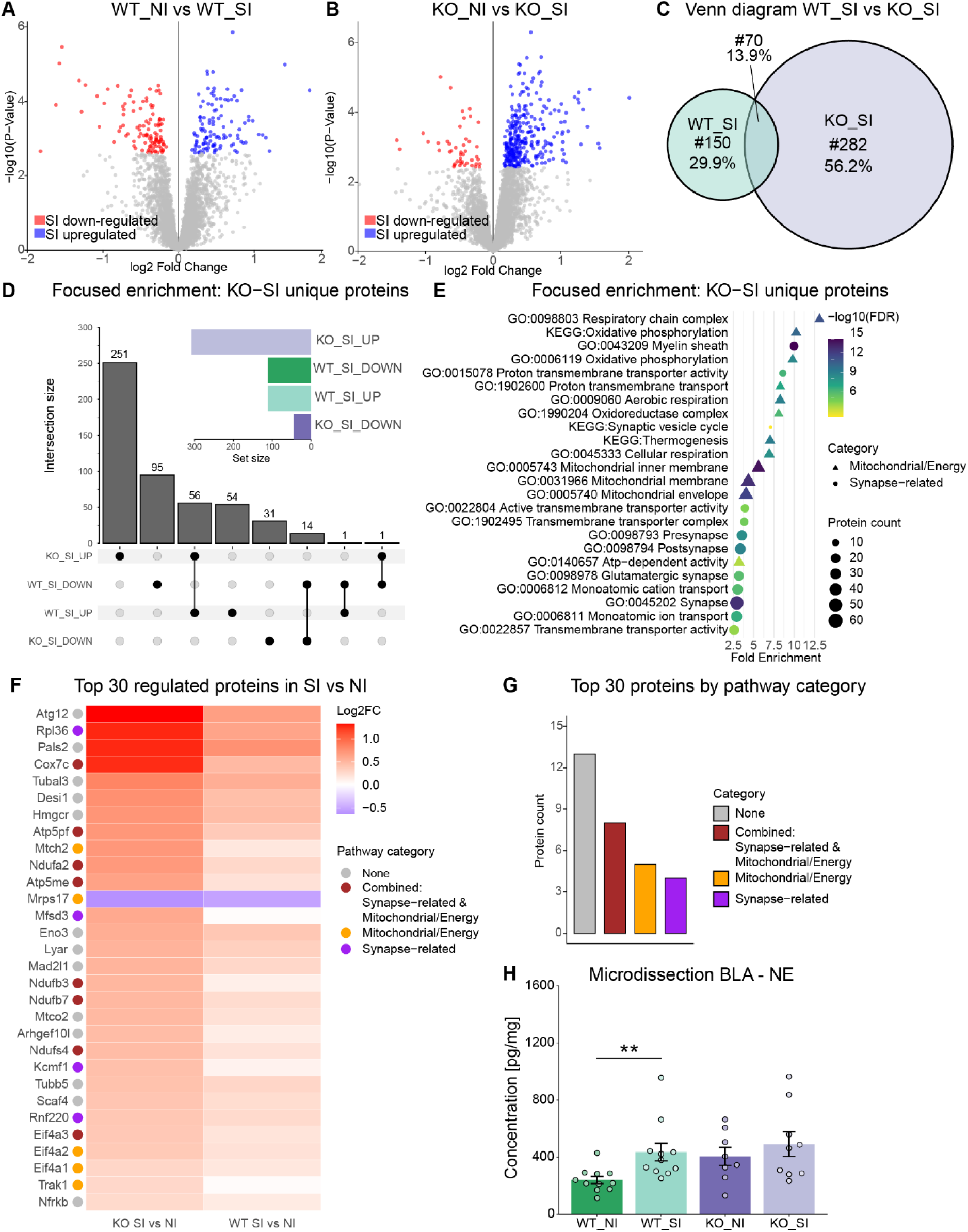
Altered mitochondrial and noradrenergic synaptic adaptations after social interaction in the BLA of *Fkbp5*^Nat^ mice. **A)** Volcano plot showing the down- and upregulated proteins in WT_NI vs WT_SI (Adj p-value<0.05: down: 111; Up: 111) **B)** Volcano plot showing the down- and up-regulated proteins in KO_NI vs KO_SI (Adj p-value<0.05: down: 46; Up: 308). **C)** Venn diagram showing the significant regulated protein overlap between WT_SI and KO_SI. **D)** UpSet plot showing the significant down- and upregulated proteins and their overlap between WT_SI and KO_SI. **E)** The proteomics GO TERM and KEGG analysis between WT and KO after NI and SI identified interesting molecular pathways related to mitochondrial, energy and synaptic structure and signaling. **F)** The top 30 regulated proteins from the “KO_SI vs KO_NI” showing a strong upregulation (29/30) of synaptic and mitochondrial/energy-related proteins. **G)** Barplot showing that from the top 30 regulated proteins, 8 proteins were annotated to both mitochondrial/energy and synaptic-related pathways, 5 were exclusively mitochondrial/energy-related, and 4 were synapse-specific. **H)** Microdissection measuring total NE levels in the BLA. Posthoc Kruskall-Wallis revealed that WT social interaction (WT_SI) significantly increased NE concentrations in the BLA compared to WT no interaction (WT_NI), F(1)=10.14, p=0.0029, which was not altered in the *Fbkp5*^Nat^ background, comparing KO_NI with KO_SI, p=0.9999. Sample size: panels A-G: WT_NI: n=10, WI_SI: n=10, KO_NI: n=11, and KO_SI n=10. Panel H WT_NI: n=11, WI_SI: n=11, KO_NI: n=8, and KO_SI n=9.

To elucidate the neurobiological mechanisms underlying the altered social behavior observed in mice lacking *Fkbp*5 and its relation to the LC-BLA circuit, a quantitative proteomic analysis was performed on the BLA. Proteomic comparisons were conducted between WT and KO mice under baseline (non-interacting, NI) conditions and immediately following a 10-minute SI task to capture the acute synaptic protein reorganization. SI induced robust and widespread proteomic changes in both genotypes when compared to NI (Figure 3A-B). Notably, the majority of regulated proteins were uniquely altered in the KO_SI group (282 proteins, 56.2%), with 150 proteins (29.9%) uniquely regulated in WT_SI, and only 70 proteins (13.9%) overlapping between genotypes (Figure 3C). Within the unique KO_SI proteome, most proteins were upregulated (251 up, 31 down), whereas in the unique WT_SI, a larger proportion were downregulated (54 up, 95 down) (Figure 3D). No direct significant genotype-dependent differences were detected under either NI (WT_NI vs. KO_NI) or SI condition (WT_SI vs. KO_SI) (Supplemental Figure 3A-B), indicating that the overall proteomic alterations induced by the KO are importantly shaped by the social experience.

To identify proteins specifically regulated in response to SI in KO mice, we focused on proteins uniquely altered (up- or downregulated) following SI (KO_SI unique; 282 proteins; Figure 3C-D). Enrichment analysis of these KO_SI–unique proteins was performed using Panther Gene Ontology (Biological Process [BP], Cellular Component [CC], and Molecular Function [MF]) and KEGG pathways. These analyses revealed a pronounced overrepresentation of mitochondrial, energy metabolism, and synapse-related pathways (Figure 3E; Supplemental Figure 3C), including oxidative phosphorylation, mitochondrial membrane organization, and synaptic vesicle cycling. Together, these findings indicate that the molecular adaptations to social experience in the BLA are tightly linked to neuronal energy metabolism and synaptic remodeling. Among the top 30 most regulated proteins in KO_SI vs KO_NI, 29 were upregulated, underscoring a strong induction profile of the KO proteome following SI, which did not reach significance in WT (Figure 3F). Several of these proteins were functionally associated with mitochondrial respiration (e.g., Cox7c, Atp5mf, Ndufa2, Mrps17) and/or synaptic structure and signaling (e.g., Tubß3, Scaf8, Eif4a1). Of these, eight proteins were annotated to both mitochondrial/energy and synaptic-related pathways, five were exclusively mitochondrial/energy-related, and 4 were synapse-specific, demonstrating that the majority of the top regulated proteins fell within these two functional categories (Figure 3G). In summary, the data indicate that *Fkbp5* deletion within the noradrenergic system leads to an altered regulation of NE availability in the BLA following SI.

### *Fkbp5*^Nat^ deletion disrupts BLA NE dynamics during social interaction

Given the observed differences in total NE levels in the BLA following SI between WT and KO, we next examined NE release dynamics during and after SI to determine whether *Fkbp5*^Nat^ deletion leads to a SI specific alteration in NE transmission within the BLA.

To assess the global real-time NE dynamics in the BLA, in-vivo microdialysis was performed before (NI) (2x 20min samples) and after a series of 20-minute SI ((4x 20min samples)) across genotypes (WT vs. KO) (Figure 4A). The MHPG/NE turnover ratio was quantified as a proxy for NE turnover and release dynamics, reflecting the balance between NE release into the extracellular space and its subsequent metabolism. In WT mice, SI led to a significant decrease in the MHPG/NE turnover rate in the BLA compared to baseline (NI), an effect not observed in KO mice (Figure 4B). A similar pattern was observed for NE concentration, which was elevated following SI relative to baseline in WT mice only (Supplemental Figure 4E). In contrast, MHPG levels remained unchanged across genotype and condition (Supplemental Figure 4F). These results indicate that the altered regulation of NE availability in KO mice is also reflected in differential NE release dynamics after SI. In order to increase the temporal understanding of the observed NE release dynamics in relation to social behavior, the genetically encoded GPCR activation-based NE sensor GRAB-NE2h^43,44^ was used to monitor extracellular NE dynamics at millisecond resolution during specific DeepOF-identified social behaviors, using in vivo miniscope imaging (Figure 4C). Viral expression and accurate targeting of the BLA were confirmed histologically (Supplemental Figure 4G-H). Analysis of global NE dynamics revealed a significant increase in NE signal when comparing the final 2 minutes of baseline NI with the first 2 minutes of the SI, in which the follow-up comparisons indicated this increase was driven by KO, with no reliable change in WT (Figure 4D-E). To examine behavior-specific NE dynamics, time-locked analysis of DeepOF social behaviors during the first 2.5 minutes of SI was performed. In WT mice, the initiation of SI, such as B-nose-to-body and B-nose-to-nose were associated with a stable Z-scored NE signal, while KO mice displayed a significant reduction in NE signal (Figure 4F-G). A similar reduction was observed during social reception, such as the W-nose-to-body interaction (Figure 4H). No differences in NE signal were observed during individual behaviors, such as B-look around (Figure 4I). It is noteworthy, that overall interaction duration times were reduced in mice implanted with the miniscope compared to non-implanted mice (Figure 2H-L vs. Supplemental Figure 5A), and no significant genotype differences in behavioral duration times were observed during the first 2.5 minutes (Supplemental Figure 5A).

**Figure 4.**
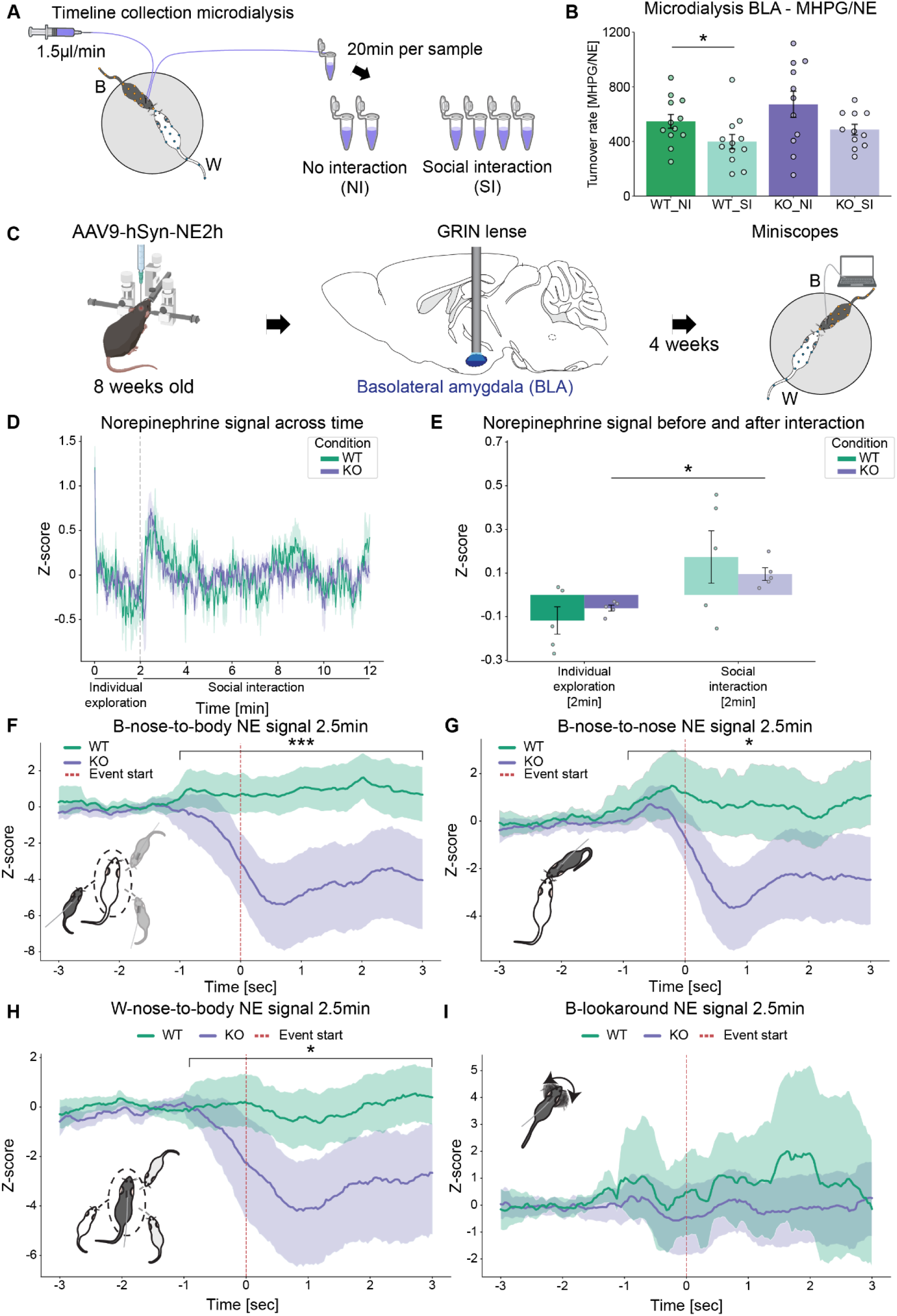
*Fkbp5*^Nat^ deletion disrupts BLA NE dynamics during social interaction. **A)** Schematic overview of the microdialysis experiment. Animals were first subjected to a 40-min baseline period, collected in two 20-min samples, followed by an 80-min social interaction (SI) phase collected across four 20-min samples. **B)** Microdialysis measuring turnover rate of MHPG/NE release in the BLA. Posthoc Tukey revealed that WT social interaction (WT_SI) has significantly lowered NE/MHPG turnover rate in the BLA compared to WT no interaction (WT_NI), F(1,22)=4.01, p=0.05, which was not altered in the *Fbkp5*^Nat^ background, comparing KO_NI with KO_SI, p=0.088. Two-way ANOVA: social interaction; F(1,42)=7.07, p=0.011, genotype; F(1,42)=0.2.87, p=0.098, social interaction*genotype; (F(1,42)=0.085, p=0.77. In panel B; n=11 for WT_NI, n=11 for WT_SI, n=8 for KO_NI, and n=9 for KO_SI. In panels C-D; n=9 for WT and n=12 for KO. In panel E; n=12 for WT and n=11 for KO (within-subject analysis on between baseline and social interaction). **C)** Schematic overview of the experimental timeline of in-vivo NE imaging using AAV9-hSyn-NE2h expressed in the BLA, visualized through a GRIN lens coupled to a miniature fluorescence microscope (miniscope). **D)** Average NE signal (Z-scored) over time during individual exploration and subsequent SI in WT and KO mice. **E)** Mean NE signal before (individual exploration) and during social interaction in WT and KO mice. Two-way ANOVA (Time × Condition) revealed a main effect of Time (NI vs SI) (F(1,16)=21.96, p=0.00025) with no effect of Condition (WT vs KO) (F(1,16)=0.20, p=0.66) and no Time × Condition interaction (F(1,16)=0.76, p=0.40). Post-hoc analysis showed a KO increase from NI to SI (p=0.012), while this was not the case comparing SI to SI under the WT condition (p=0.240). **F-I)** Event-aligned NE signal traces (±3 s) around specific DeepOF identified social behaviors; **F**) The NE signal during B–nose-to-body in KO mice showed a significant reduction compared to WT (Mixed linear model β₍KO₎ = −522.04 ± 134.73, z = −3.88, p < 0.001). **G)** The NE signal during B–nose-to-nose in KO mice showed a significant reduction compared to WT (Mixed linear model β₍KO₎ = −313.69 ± 149.02, z = −2.11, p = 0.035). **H)** The NE signal during W–nose-to-body in KO mice showed a significant reduction compared to WT (Mixed linear model β₍KO₎ = −318.96 ± 140.67, z = −2.27, p = 0.023). **I)** The NE signal during B–lookaround in KO mice did not differ from WT (Mixed linear model β₍KO₎ = −115.51 ± 147.75, z = −0.78, p = 0.434). Sample size: panels A-B: WT: n=12 and KO n=11. Panels C-I WT: n=5 and KO n=5.

To evaluate the persistence of these effects, NE dynamics were analyzed across the full 10-minute SI session. The initiation of SI, via B-nose-to-body and the social reception, via W-nose-to-body, continued to exhibit a stable NE signal in WT mice, while KO mice showed a significant reduction (Supplemental Figure 5B-D). However, no genotype effect was observed during B-nose-to-nose interaction over the 10min time bin data (Supplemental Figure 5C). As in the 2.5min time window, individual behaviors, including B-look around remained unaffected by genotype (Supplemental Figure 5E). Interaction duration times over the full 10min were comparable to those seen in the first 2.5min of non-implanted animals (Figure 2H-L & Supplemental Figure 5F), and no genotype-related differences in time duration were observed for any social and individual behaviors (Supplemental Figure 5F). Together, these findings demonstrate that *Fkbp5*^Nat^ deletion leads to a specific reduction in BLA NE signaling during active social engagement, while NE signaling during individual, non-social behaviors remain unaffected.

### *Fkbp5*^Nat^ deletion is not affecting BLA NE Dynamics During Social Interaction with same strain C57Bl/6N

To assess whether the social behavior patterns, observed during SI with an unfamiliar CD1 mouse, were specific to that context, the DeepOF behaviors were examined during interaction with an unfamiliar conspecific of the same strain (C57BL/6N). In contrast to the CD1 interaction, KO mice displayed a significant reduction in SI duration during the first 2.5 minutes, specifically in B^test^-nose-to-body, with trends toward reduced duration in B^test^-nose-to-nose and B^stim^-nose-to-body (Figure 5A). Non-social behaviors, including B^test^-look around, showed no genotype-related differences. These behavioral differences diminished over time, consistent with the habituation effects observed in non-implanted mice during CD1 interaction (Supplemental Figure 2A-B). When considering the full 10-minute session, no significant genotype differences remained in any of the measured behaviors, including social and individual behaviors (Supplemental Figure 6A).

**Figure 5.**
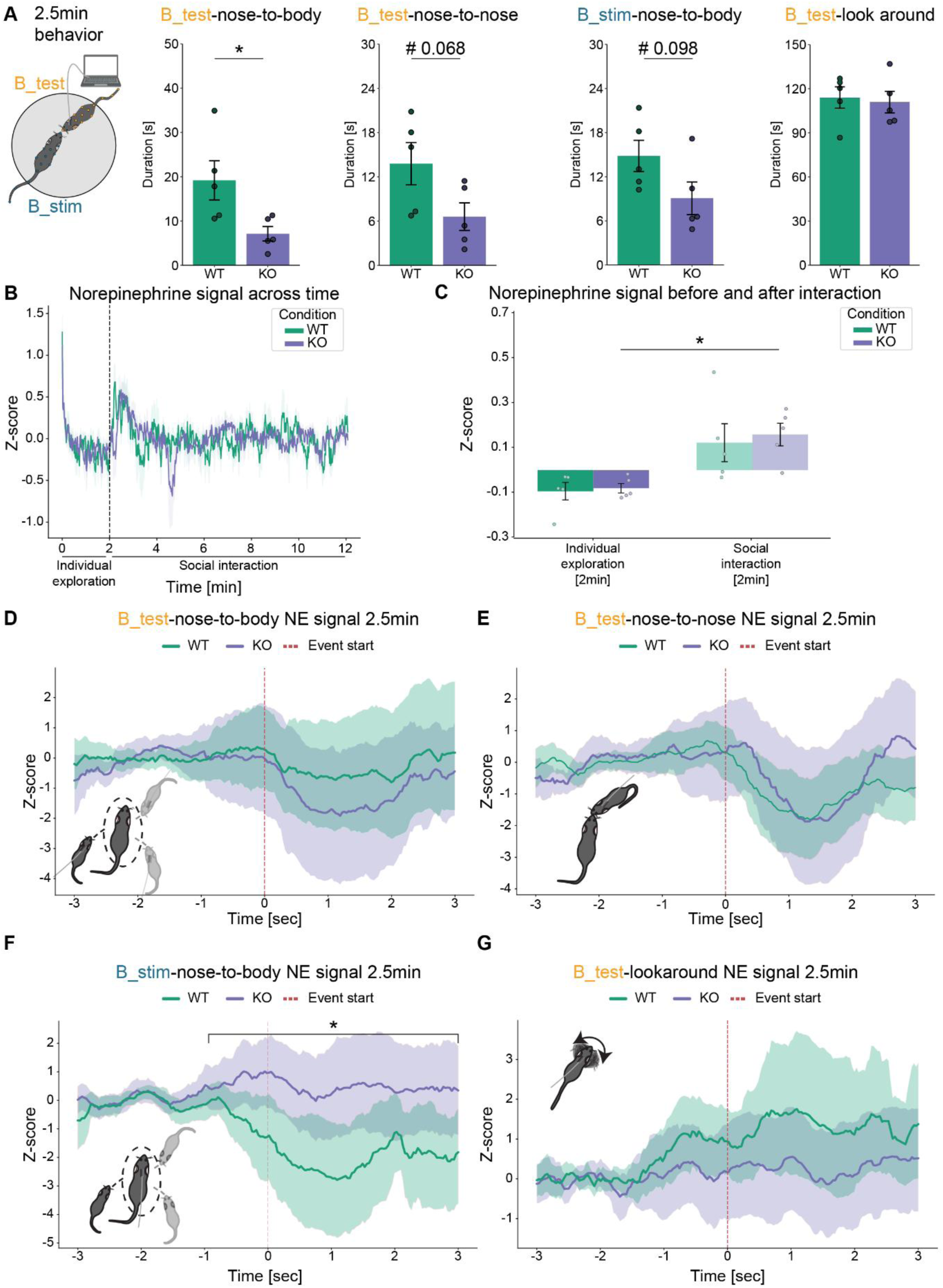
*Fkbp5*^Nat^ deletion does not disrupts BLA NE dynamics during same strain C57/Bl6N social interaction. **A)** Schematic overview of the test animal (B_test) connected to in-vivo NE imaging during social interaction with a same strain conspecific (B_stim). A significant decrease in KO compared to WT mice for B_test-nose-to-body was observed (T(8) = −2.56, p = 0.0336), while a trend to a decrease in KO compared to WT was observed for B_test-nose-to-nose (T(8) = −2.10, p = 0.0686) and B_stim-nose-to-body (T(8) = −1.87, p = 0.0978). No significant difference between WT and KO was observed in the B_test-look around behavior (T(8) = −0.29, p = 0.7780). **B)** Average NE signal (Z-scored) over time during individual exploration and subsequent SI with same strain C57/Bl6N in WT and KO mice. **C)** Mean NE signal before (individual exploration) and during social interaction in WT and KO mice. Two-way ANOVA (Time × Condition) revealed a main effect of Time (NI vs SI) (F(1,16)=19.73, p=0.00041) with no effect of Condition (WT vs KO) (F(1,16)=0.15, p=0.705) and no Time × Condition interaction (F(1,16)=0.04, p=0.848). Post-hoc analysis showed a KO increase from NI to SI (p=0.0229), while this was not the case comparing NI to SI under the WT condition (p=0.200). **D-G)** Event-aligned NE signal traces (±3 s) around specific DeepOF identified social behaviors; **D**) The NE signal during B_test–nose-to-body in KO mice did not differ from WT (Mixed linear model β₍KO₎ = −86.42 ± 187.67, z = −0.46, p = 0.645). **E)** The NE signal during B_test–nose-to-nose in KO mice did not differ from WT (Mixed linear model β₍KO₎ = 54.91 ± 130.18, z = 0.42, p = 0.673). **F)** The NE signal during B_stim–nose-to-body in KO mice showed a significant increase compared to WT (Mixed linear model β₍KO₎ = 261.65 ± 112.72, z = 2.32, p = 0.020). **G)** The NE signal during B_test–lookaround in KO mice did not differ from WT (Mixed linear model β₍KO₎ = −114.43 ± 105.09, z = −1.09, p = 0.276). Sample size: panels A-E: WT: n=5 and KO n=5.

Analysis of global NE dynamics during interaction with unfamiliar C57BL/6N conspecifics revealed a significant increase in NE signal from the final 2 minutes of NI to the first 2 minutes of SI; follow-up comparisons again indicated this increase was driven by KO, with no reliable change in WT, mirroring the CD1 interaction data (Figure 5B-C). However, the 2.5min bin time-locked analysis of NE dynamics during interaction with C57BL/6N conspecifics revealed, in contrast to the CD1 interaction findings, no significant genotype differences in NE signal during the initiation of SI (B^test^-nose-to-body and B^test^-nose-to-nose; Figure 5D-E). Furthermore, compared to the CD1 social reception findings (W-nose-to-body; Figure 4F), the B^stim^-nose-to-body social reception showed an opposite pattern, with WT mice displaying a significantly lower NE signal than KO mice (Figure 5F). As with CD1 interactions, no genotype differences in NE dynamics were observed during non-social behaviors such as B-look around (Figure 5G). Extending the time-locked analysis to the full 10-minute SI confirmed the absence of genotype differences across social initiation (B^test^-nose-to-body, B^test^-nose-to-nose; Supplemental Figure 6 B-C), social reception (B^stim^-nose-to-body; Supplemental Figure 6D), and non-social behaviors (B^test^-look around; Supplemental Figure 6E). These results indicate that the NE signaling differences associated with *Fkbp5*^Nat^ deletion are context-dependent and graded, becoming more pronounced as the conspecific novelty increases.

## Discussion

The LC is a crucial node within the stress response system^32^, and a combination of severe stress exposure and genetic risk variants of the *FKBP5* gene have been found to increase the risk for MDD pathology^22,45,46^. The current study shows the selectivity of the LC-BLA circuitry towards social stressors through cellular activation and *Fkbp5* signaling, thereby sex-specifically shaping social behavior in a contextually scalable manner.

### *Fkbp5* expression in the LC reacts to stress in a highly selective way

Previous studies have provided evidence supporting the brain region-specific upregulation of *Fkbp5* mRNA, which is contingent upon the specific type of stressor employed^24–26^. The present study uncovers a highly selective upregulation of *Fkbp5* in the LC following acute social stress but not acute restraint stress or acute social stress after a background of chronic social stress. Interestingly, the DRN region did not follow the same *Fkbp5* regulation pattern, as we observed that both acute social stress only and acute social stress after a background of chronic social stress led to an upregulation of *Fkbp5* in the DRN. Consistent with previous research demonstrating brain-region-specific upregulation of *Fkbp5*^24^, this study shows that the regulation of *Fkbp5* expression in response to social stress varies across different brain regions. Previous research has shown that acute stress exposure differentially alters LC excitability over time, with increased LC excitability observed one week after stress exposure compared to immediately after the stress exposure^47^. The specific increase in *Fkbp5* mRNA levels may be attributed to heightened cellular activity in the LC region during acute social stress. To exclude the possibility that the observed differential expression pattern of *Fkbp5* is merely a reflection of cellular activity, we examined the expression c-Fos as an indicator of cellular activity^48–51^ across the different stress groups. Interestingly, c-Fos protein expression levels in the LC induced a robust c-Fos upregulation, which was not altered compared to acute social stress after a history of chronic social defeat stress and thereby different from the observed *Fkbp5* expression pattern. This finding aligns with Reyes et al., 2019^52^, who demonstrated that a shorter 5-day CSDS protocol exhibited a much lower c-Fos count compared to acute social stress. Furthermore, Torres-Sánchez et al., 2025^53^ showed that ventromedial prefrontal cortex DBS switches LC responses to social defeat from activation to inhibition and stabilizes auditory-evoked LC responses. Accordingly, the differential LC *Fkbp5* expression we observe across stress history is unlikely to be explained solely by overall LC firing. In the LC, *Fkbp5* is robustly upregulated after acute social stress but attenuated after repeated exposure, suggesting regulation by the social-stress context and positioning LC *Fkbp5* as a promising therapeutic target to modulate specifically social behavior.

### Noradrenergic *Fkbp5* deletion alters the social behavioral profile in male, but not female mice

Considering the high selectivity of the *Fkbp5* LC regulation after social stress, we tested whether the lack of *Fkbp5* within the noradrenergic system, including the LC, shapes social behavior. The behavioral profile was investigated using advanced automated behavioral assessment tools^41^ in freely moving mouse dyads, thereby fully capturing the complexity of the social behavior^38^. The overall social behavioral profile was selectively altered in male KO mice, indicating pro-social behavior. More specifically, KO males showed increased nose-to-tail, nose-to-body, and following, whereas in previous work we showed that chronic social stress produced the opposite pattern, reducing nose-to-tail and nose-to-body behaviors, pointing to a selective shift toward social approach^41^. In contrast, in female mice or after male chronic social stress exposure, no changes in the social behavioral profile were observed. Thus, the behavioral effects map onto the LC *Fkbp5* regulation pattern: changes emerge with acute social exposure and are attenuated after chronic stress. Together, our findings indicate that noradrenergic *Fkbp5* gates social approach behavior in a sex- and stress-history dependent manner, with effects expressed in males during ethological, acute social encounters and lost after repeated stress. These data therefore show LC-linked *Fkbp5* as a valid target to dissect how genetic risk, sex, and social context interact to shape social behavior, and highlight it as a promising target for interventions aiming to normalize social salience without globally suppressing noradrenergic function.

### Noradrenergic *Fkbp5* deletion alters the mitochondrial and synaptic pathways after social interaction in male mice

Our data establish the BLA as a central node through which *Fkbp5* regulates SI circuits. *Fkbp5* deletion selectively abolished the SI-evoked increase in NE output in the BLA, but not in other major locus coeruleus targets. This key circuit modulation is likely mediated by an underlying alteration in synaptic infrastructure, as supported by BLA proteomic profiling. Following SI, *Fkbp5* KO mice showed a pronounced, genotype-specific proteomic shift, with significant enrichment in pathways related to mitochondrial function and synapse organization, indicating that *Fkbp5* is central for maintaining the noradrenergic synaptic competence required for social behavioral modulation. In line with our findings, Sandi and others have repeatedly shown that chronic stress prominently alters mitochondria-associated gene pathways, with enrichment for oxidative phosphorylation and related energy processes across different brain regions^54^, which can directly impact stress-related behaviors, including anxiety^55^. Interestingly, they further show that manipulating mitochondrial function in the nucleus accumbens alters social dominance and energy metabolism^56,57^. These findings align with our BLA proteomics results, where KO_SI exhibits selective enrichment of mitochondrial/energy and synapse-related pathways alongside the loss of social-evoked NE mobilization in KO males. The increased energy and mitochondrial workload in KO_SI could be mediated through the autophagy pathway, since our findings show that the key modulator of autophagy, Atg12, is most strongly regulated during KO_SI. Interestingly, previous research has shown a directly interaction between *Fkbp5* and Atg12, and have linked this to disease phenotype and antidepressant treatment response^23,58,59^. Together, in line with this prior work, our data provide evidence that *Fkbp5*-dependent noradrenergic signaling interfaces with mitochondrial state in the BLA to shape social interaction responses, rather than reflecting a nonspecific stress effect.

### Noradrenergic *Fkbp5* deletion disrupts BLA NE dynamics during social interaction in a context-dependent manner

To dissect the context-specific nature of *Fkbp5*’s role in the noradrenergic system, we utilized a SI paradigm comparing outbred (CD1) versus same-strain (C57BL/6N) conspecifics. We found that the *Fkbp5* deletion selectively impaired NE mobilization in the BLA only when interacting with a high-salience, outbred CD1 mouse. Specifically, microdialysis confirmed the loss of the typical NE surge, and miniscope recording revealed a significant reduction in phasic, behavior-locked NE transients during both active social engagement and social reception in KO mice, suggesting a failure to appropriately encode novel social stimuli. In contrast, when interacting with a lower-salience, same-strain C57BL/6N mouse, the BLA NE dynamics were largely preserved or even reversed between genotypes. Taken together, these findings suggest that *Fkbp5* is crucial for modulating BLA NE output specifically in response to socially salient or novel stimuli. The preservation or reversal of NE dynamics during C57BL/6N interaction, coupled with the KO mice showing less interest in C57BL/6N conspecifics (a behavioral reversal relative to the main findings), highlights that *Fkbp5* dictates how the BLA assesses and responds to social salience across different interaction contexts. Previous research has established the crucial role of the noradrenergic pathway in the BLA, where it is heavily involved in emotional memory consolidation^60–62^. Additionally, McCall et al. 2017^32^ demonstrated that specific stimulation of LC-NE fibers in the BLA leads to NE release, alters BLA activity patterns, and increases anxiety-like behavior. These studies demonstrate the general importance of the LC-BLA pathway for broad behavioral and physiological regulation, and our current data represent a critical advancement by revealing the *Fkbp5*-mediated mechanism that specifically governs social salience encoding via context-dependent noradrenergic output in the BLA. The convergence of the data supports a state-dependent gating model in which *Fkbp5* in noradrenergic neurons calibrates phasic NE transients during stronger socially salient events. The unaffected, or even reversed, NE responses during same-strain encounters suggest that when social salience is lower or more familiar, compensatory mechanisms maintain BLA NE transmission despite *Fkbp5* deletion. This is consistent with the reported role of the BLA as a central hub for generating and encoding emotional salience signals, often exhibiting graded responses to familiar versus novel cues^63,64^. This state-dependent gating model is mechanistically underpinned by our BLA proteomics data, which revealed a substantial *Fkbp5*-specific remodeling of mitochondrial/energy and synapse-related pathways following social interaction. We propose that the deletion of *Fkbp5* compromises the metabolic-synaptic coupling essential for generating the high-gain, phasic NE release necessary to encode highly salient social events, thereby leaving baseline or low-salience transmission relatively preserved.

## Online Methods

### Animals and housing

Wild-type adult male and female C57/Bl6N mice and the genetic mouse line *Fkbp5*^Nat^ (age between 2-3 months) were obtained from the in-house breeding facility of the Max Planck Institute of Psychiatry and used for breeding (F_0_). The offspring (F_1_) were used as experimental animals and were weaned at P25 in groups of maximum four animals with same-sex littermates. All experiments were performed during adulthood at the age of 3-5 months. All animals were housed in individually-ventilated cages (IVC; 30cm×16cm×16cm connected by a central airflow system: Tecniplast, IVC Green Line—GM500), while kept under standard housing conditions; 12h/12h light-dark cycle (lights on at 7 a.m.), temperature 23±1°C, humidity 55%. Food (Altromin 1324, Altromin GmbH, Germany) and water were available ad libitum. The experimental procedures were approved by the committee for the Care and Use of Laboratory Animals of the government of Upper Bavaria, Germany. All experiments were in accordance with the European Communities Council Directive 2010/63/EU.

### Generation of the *Fkbp5*^Nat^ mouse line

The genetic mouse line *Fkbp5*^Nat^ is a conditional knockout of the *Fkbp5* gene in the noradrenergic system, among others, the LC. This was achieved via the generation (as performed by the knockout mouse project) of full knockout *Fkbp5*^Frt/Frt^ mice, which can re-express functional *Fkbp5* upon activation by the Flp recombinase. *Fkbp5*^Frt/Frt^ mice were bred with Deleter-Flpe mice to create mice with a floxed *Fkbp5* gene; *Fkbp5*^lox/lox^ mice^65,66^. Subsequently, the final conditional knockout of *Fkbp5* was achieved via the crossing of *Fkbp5*^lox/lox^ mice with a BAC transgenic mouse line expressing Cre recombinase under the control of the noradrenalin transporter gene (Nat, Tg(Slc6a2-cre)FV319Gsat)^67^, resulting in *Fkbp5*^Nat^ mice. The *Fkbp5*^lox/lox^ litter mates were used as wild-type (WT) control animals.

### Chronic social defeat stress

A cohort of wild-type C57/Bl6N 2 months-old male mice was either subjected to the chronic social defeat stress (CSDS) protocol or was kept under normal housing conditions. The CSDS paradigm consisted of exposing the experimental C57Bl/6N mice to an aggressive CD1 mouse for 21 consecutive days, as previously described^68^. The CD1 aggressor male mice were purchased from Janvier Labs (Germany) and were at least 16 weeks old. The CD1 aggressor mice were trained and selected based on their aggression prior to the start of the experiment. Upon the start of the CSDS, the experimental mice were introduced daily to a new CD1 resident’s territory, who subsequently attacked and forced the experimental mouse into subordination. Defeat sessions lasted until the stress-exposed mouse received two bouts of attacks from the CD1 aggressor, or at five minutes in the rare instances when two bouts were not achieved within this duration. Between daily defeats, stressed mice were housed in the resident’s home-cage but physically separated from the resident by a see-through, perforated mesh barrier, allowing sensory exposure to the CD1 aggressor mouse while preventing further attacks. The defeat time of day was randomized between 11 a.m. and 6 p.m. to avoid habituation and anticipatory behaviors in defeated mice. NS mice were single-housed in the same room as the stressed mice. All animals were handled daily and weighed every 3-4 days.

### Acute stress exposure

The acute stress exposure consisted of either acute social defeat stress or restraint stress. The acute stress was performed 24 hours after the end of the CSDS paradigm. The acute social defeat stress consisted of a single defeat event with a novel CD1 aggressor mouse, as described for the CSDS paradigm. The restraint stress consisted of restraining in a 50ml falcon tube, which contained holes in the top and the lid to allow air ventilation and tail movement. Animals were restrained for 15min during which they were kept in their home-cage environment. Animals were euthanized 4 hours after acute stress exposure, after which, brains were removed and snap-frozen using 2-methyl butane (kept on dry ice) and stored at −80°C until further use.

### *In-situ* hybridization of *Fkbp5*

The *Fkbp5* mRNA profile was determined using radioactive *in-situ* hybridization labeling as described previously^24^. In brief, brains were sliced using a cryostat in 20 µm sagittal sections, which resulted in a series of LC (bregma: −5.34 to −5.80) and DRN (bregma: −4.36 to −4.84) slides that were thaw-mounted on Super Frost Plus Slides and stored at −20°C. The *in-situ* hybridization sections were removed from −20°C, left to dry at room temperature, fixated with 4% paraformaldehyde, and subsequently dehydrated using a series of increasing concentrations of ethanol. Then, the hybridization buffer was equally spread out over the different slides containing the radioactive ^35^S-UTP-labeled *Fkbp5* riboprobe and incubated overnight at 55°C. The next day, the sections were rinsed, incubated with RNAse A, desalted, and dehydrated, after which the radioactive slides were exposed to Kodak Biomax MR films (Eastman Kodak Co., Rochester, NY) and developed after an exposure time of 12 days. Films were digitized, and the regions of interest were identified using the mouse brain atlas (https://mouse.brain-map.org/). The expression was determined by optical densitometry with the ImageJ software (NIH, Bethesda, MD, USA). The expression was averaged per brain region per animal and subtracted by the background signal of a nearby structure that did not express the *Fkbp5* gene.

### Immunostaining of c-Fos

The c-Fos protein expression was determined using a DAB staining kit for immunohistochemistry (Abcam, USA; ab64261). Brains were sliced using a cryostat in 20 µm sagittal sections, which resulted in a series of LC slides that were thaw-mounted on Super Frost Plus Slides and stored at −20°C. Immunostaining was performed as described by the Abcam DAB staining protocol (ab64261). In short, the slides were fixated in 4% paraformaldehyde, then incubated with hydrogen peroxide (to block endogenous peroxidase), and then blocked in protein block to minimize unspecific binding. Then slides were incubated overnight at 4°C with the rabbit monoclonal c-Fos primary antibody (1:1000, ab222699), diluted in phosphate buffered saline (PBS) and 0.5% Bovine Serum Albumin. On the next day, the sections were incubated at room temperature for 10 min with the secondary antibody goat anti-polyvalent. Next, to amplify the signal, slides were incubated with streptavidin peroxidase. The DAB staining was performed by combining the DAB chromogen with the DAB substrate (1:50, respectively), which was then applied to the slides and exactly washed away after 3.5 min. Slides were dehydrated using a series of increasing concentrations of ethanol and cover slipped. Slides were then imaged using a slide scanner (Olympus, VS120-S6-W) on a 10x magnification using the bright field settings. Bilateral images were taken in a series of images of the LC, going from bregma −5.34 to −5.80. Ultimately, similar bregma images were taken for all animals, and c-Fos puncta were counted for 2-3 images per animal per side.

### Validation of the *Fkbp5*^Nat^ mouse line using RNAscope

Validation of the knockout of *Fkbp5* in the LC was performed via an RNAscope *in-situ* hybridization study. Male mice were euthanized under baseline conditions at 3-5 months of age. The brains were removed and snap-frozen using 2-methyl butane (kept on dry ice) and stored at −80°C until further use. Brains were sliced using a cryostat in 20 µm sagittal sections, which resulted in a series of LC slides that were thaw-mounted on Super Frost Plus Slides and stored at −20°C. The RNAscope staining procedures were used according to the manufacturer’s protocol as previously described^66^. The RNAscope fluorescent multiplex reagent kit (cat. no. 320850, Advanced Cell Diagnostics, Newark, CA, USA) was utilized for mRNA staining. The probes used for staining were *Fkbp5* (Probe: Mm-*Fkbp*5-C1), and Tyrosine hydroxylase (TH) (Probe: Mm-TH-C2). Slides were then imaged using a ZEISS confocal microscope on a 40x magnification using the fluorescent channel. Bilateral images were taken in a series of images of the LC, going from bregma −5.34 to −5.52. All images were acquired using the same settings for laser power, detector gain, and amplifier offset. *Fkbp*5 mRNA expression was analyzed using ImageJ, with the experimenter blinded to the genotype of the animals, and was counted manually. Similar bregma images were taken for all animals and *Fkbp5* puncta were counted within TH-positive cells for 2-3 images per animal per side. *Fkbp5* negative TH cells were accounted for when the cell had less than five *Fkbp5* puncta. Ultimately, a calculation per animal was made for the percentage of *Fkbp5-positive* TH cells compared to the total amount of TH cells.

### *Fkbp5*^Nat^ adult testing

At 3 months of age, a cohort of nonstressed males and females was tested in the social interaction (SI) task. In addition, separate cohorts of male *Fkbp5*^Nat^ were used to do microdissection, proteomics, microdialysis, and miniscope experiments. The behavioral tests were performed between 8 a.m. and 11 a.m. in the same room as the housing facility.

### Social interaction task analysis using DeepOF

The SI task was performed in a round open field arena (diameter of 38cm) using sawdust material at the bottom, as previously described by Bordes et al^41,42^. The experimental animal was placed in the open field arena and could freely explore for 10 min, after which an unfamiliar young CD1 (4–6 weeks old) social conspecific was placed in the same arena and both were allowed free exploration of the arena and each other for 10min. The data was recorded using the DFK37BUX250 imaging source (Germany) cameras with camera lenses from Stoelting, Ireland (Item nr. 60528). The IC capture software (version 2.5.1547.4007) from imaging source was used to obtain the videos, and further analysis was performed with DeepLabCut version 2.2b7 (single animal mode for pose estimation^39,40^), and subsequently, DeepOF module version 0.4.6^42^ for supervised behavioral analysis of six individualistic behaviors during the open field; wall-climbing, digging, huddling, look-around, sniffing, and speed (locomotion), and all during the SI task, including an additional five social behaviors; nose-to-nose, Side-by-side, Side-reverse-side, nose-to-tail, and Nose-to-body. Data were analyzed for the total 10 min of both tasks and in time bins of 2.5 min.

### Microdissection

The total concentration of NE was measured from microdissected fresh frozen brain tissue in both WT and KO mice, comparing no interaction (NI) and SI conditions. At the start of the experiment, mice were randomly divided into either the NI or SI condition. The NI animals were left undisturbed in their home-cage, whereas SI animals were exposed to an unfamiliar young CD1 (4–6 weeks old) mouse for 10min within their home-cage. After the 10min SI exposure, all animals were directly euthanized, and brains were removed and snap-frozen using 2-methyl butane (kept on dry ice) and stored at −80°C until further use. Then, the brains were sectioned using a VT1200/S Leica vibratome on 250µm thick slices by the different brain regions; 2 slices in the medial prefrontal cortex (mPFC) (bregma: 1.94 to 1.54), 3 slices in the basolateral amygdala (BLA) (bregma: −1.34 to −1.94), 3 slices in the dorsal hippocampus (dHipp) (bregma: −1.70 to-2.18), and 3 slices in the ventral hippocampus (bregma: −3.08 to −3.52). The sliced tissue was directly punched within the vibratome using a sample corer (diameter 1 mm) and stored in 1.5 mL Safe-lock Eppendorf tubes on dry ice and subsequently stored at −80°C until further use. The measurement of NE was carried out by reverse-phase liquid chromatography with electrochemical detection as described in Nagler et al., 2018^69^. The values obtained were expressed as picograms per milligram (pg/mg) wet tissue and were logarithmically transformed for calculation of linearity of regression, standard error of the regression coefficients, and significance of differences between regression coefficients.

### Proteomics

The proteomics analysis was performed from BLA microdissected fresh frozen brain tissue in both WT and KO mice after NI and after the 10 min SI. The same behavioral and extraction protocol was performed as described during the microdissection. BLA tissue punches were homogenized in ice-cold T-PER™ tissue protein extraction reagent (ThermoFisher Scientific, 78510) freshly supplemented with protease inhibitor cocktail tablets (Roche, 05892791001) and phosphatase inhibitor cocktail tablets (Roche, 04906837001). Protein extracts were centrifuged at 10,000 x g for 10 minutes to pellet cell/tissue debris. Subsequently, protein concentration was adjusted to 2µg/µl using the Pierce™ BCA protein assay kit (ThermoFisher Scientific, 23225). For mass spectrometry measurements, in-solution samples were sent to the Max Planck Institute of Biochemistry Core Facility, Mass Spectrometry Lab.

### Protein Aggregation Capture (PAC) Sample Preparation, Digestion and Phospho-Peptide Enrichment

Magnetic beads (Resyn magnetic beads, Hydroxyl modified, 20 µg/µl stock) were prepared in a 2:1 ratio relative to the estimated amount of protein, using 10 µL of bead suspension per 100 µg of protein lysate (bead stock concentration: 20 µg/µL). Beads were washed two times with 70% (v/v) acetonitrile (ACN) to remove storage buffer and for equilibration. Each protein sample was diluted to a final volume of 350 µL with 70% (v/v) acetonitrile (ACN), after which the pre-washed beads were added. Samples were vortexed for approximately 10 s and subsequently incubated on ice for 10 min without agitation. This vortexing and incubation step was repeated once more to ensure efficient protein aggregation and capture on the beads. Beads were pelleted using a magnetic rack, and the supernatant was carefully removed. The beads were sequentially washed with 100% ACN, 95% ACN, and 70% ethanol (EtOH). After the final wash, residual 70% EtOH was removed, and the beads were allowed to air-dry for approximately 5 min, ensuring that they did not dry completely.

Beads were resuspended in 50 µL of digestion buffer (50 mM TEAB pH 8.5). Samples were incubated at room temperature for 30 min with continuous shaking. Digestion was initiated by the addition of sequencing-grade modified trypsin and LysC at enzyme-to-protein ratios of 1:100 and 1:200 (w/w), respectively. Samples were incubated overnight at 37 °C with shaking at 1100 rpm. The digestion was quenched by adding trifluoroacetic acid (TFA) to a final concentration of 1% (v/v) followed by desalting of the peptide mixture via Sep-Pak C18 1cc vacuum cartridges^70^. Phosphorylated peptides were enriched with Fe (III)-NTA cartridges (Agilent Technologies; Santa Clara, Ca) using the AssayMAP Bravo Platform (Agilent Technologies; Santa Clara, Ca) according to published protocols^71^. Approximately 200ng of non-enriched and enriched samples were loaded onto Evotips.

### LC-MS/MS data acquisition

Peptides were eluted from the Evotips onto a 15 cm PepSep C18 column (1.5 µm, Bruker Daltonics) using the Evosep One HPLC system (Evosep). The column was maintained at 50 °C, and peptide separation was achieved using the 30 SPD method. Eluted peptides were directly ionized and introduced into a timsTOF Pro mass spectrometer (Bruker) via electrospray ionization. Data acquisition was performed in data-independent acquisition (DIA) PASEF mode via timsControl. Mass spectrometry covered a scan range of 100–1700 m/z, and ion mobility ranged from 1/K0 = 0.70 to 1.30 Vs·cm². The dual TIMS analyzer utilized equal ion accumulation and ramp times of 100 ms each, with a spectra rate of 9.52 Hz. For DIA-PASEF scans, the mass scan range was 350.2–1199.9 Da, and ion mobility ranged from 1/K0 = 0.70 to 1.30 Vs·cm². Collision energy was linearly ramped based on ion mobility, from 45 eV at 1/K0 = 1.30 Vs·cm² to 27 eV at 1/K0 = 0.85 Vs·cm². A total of 42 DIA-PASEF windows were acquired per TIMS scan, with switching precursor isolation windows, resulting in an estimated cycle time of 2.21 seconds.

### Data analysis

The raw data were analyzed using Spectronaut 19.0 in directDIA+ (library-free) mode, applying standard settings. The peak list was compared to a predicted library from the Mouse UniProt database (downloaded in 2023, SwissProt entries). Cysteine carbamidomethylation was set as a fixed modification, while methionine oxidation, N-terminal acetylation and phosphorylation (on S, T, Y) were used as variable modifications. Quantification was performed across samples using label-free quantification (MaxLFQ) at the MS2 level.

Bioinformatic analyses were conducted using R (v4.3.2). For phosphoproteomics, only phosphosites with a localization probability greater than 0.75 were used for analysis and one outlier sample identified during QC was removed. For proteomic and phosphoproteomic analysis, sites/proteins with ≥70% observed values across samples were kept and intensities were log2-transformed. Differential abundance was assessed with moderated linear models (limma) using the factors Genotype (KO vs WT) and NI (SI vs NI). Significance was controlled at FDR < 0.05 (Benjamini–Hochberg); effect sizes are reported as log2 fold changes. Analyses were performed in R using limma.

### In-vivo microdialysis

The microdialysis experiment consisted of the measurement of NE and its metabolite 3-Methoxy-4-hydroxyphenylglycol (MHPG) unilaterally in the right BLA during NI and SI in WT and KO animals. The microdialysis workflow was performed as described previously^72^. In brief, male mice at 3 months of age were anesthetized with isoflurane and fixated in a stereotactic apparatus. Then microdialysis guide cannula was inserted in the right BLA (bregma: AP −1.35mm, ML 3.3 mm, and DV 4.4mm). After surgery, animals were treated with meloxicam for three days and were allowed to recover for a minimum of 7 days in the home-cage environment.

The microdialysis system contained a syringe pump (Harvard Apparatus, USA) that was connected via FEP tubing (CMA, Cat. N. 8409501) to a dual-channel liquid swivel (Microbiotech Se) and could then be connected to the probe via FEB tubing. The perfusion liquid was artificial CSF, which consisted of a solution of NaCl (0.86%), KCl (0.020%), MgClx6H_2_O (0.024%), and CaClx2H_2_O (0.018%) in HPLC water with a PH of 7.4. Then, the microdialysis probe (CMA 7 Probe 1 mm membrane, 6 kDa; Cat.N. 000082) was connected to the running microdialysis system using a constant flow rate of 1.5µl/min. Any air bubbles in the probe were removed, after which the probe was inserted by hand into the guide cannula of the animal at least 20 hours before the start of the experiment. Animals stayed in specific microdialysis cages containing a standard amount of sawdust material and nesting material (16 cm length x 16 cm width x 32 cm height). The experiment started with the collection of baseline samples, after which an unfamiliar young CD1 (4–6 weeks old) mouse was introduced into the microdialysis cages, resulting in the comparison of 2x 20min baseline, NI samples with 4x 20min SI samples. The measurement of NE and MHPG out of the microdialysates was performed using HPLC with electrical detection, as previously described by Anderzhanova et al., 2020^72^. Quantification was performed using external standard calibration (0.1–5 nM).

### In-vivo analysis of norepinephrine signaling using miniscopes

#### Surgical procedures

Mice (2 months old) were anesthetized with isoflurane and secured in a stereotactic frame. A total volume of 500 nL of AAV9-hSyn-NE2h (WZ Biosciences, USA)^43,44^ was injected at a rate of 100 nL/min into the right basolateral amygdala (BLA; coordinates: AP = –1.35, ML = 3.3, DV = 4.7 mm) to express the genetically encoded norepinephrine (NE) sensor GRAB-NE2h in neurons. Post-surgery, animals received meloxicam for three consecutive days.

After at least 3 weeks of recovery, a gradient-index (GRIN) lens was implanted. A small craniotomy was created by slowly lowering a 20 G needle (with the tip cut) into the brain to a depth 400 µm dorsal to the viral injection site (DV = 4.3 mm), to clear the path for the lens. The needle was then removed and a GRIN lens (ProView, Inscopix; diameter: 0.6 mm, length: ∼7.3 mm) was slowly lowered to a final position 200 µm dorsal to the injection site (DV = 4.5 mm). The skull was first sealed with Histoacryl glue (Braun), and the lens was then secured using dental acrylic.

Following lens implantation, animals were allowed to recover for at least 6 weeks. Then, a baseplate (BPL-2, Inscopix) was mounted above the GRIN lens, adjusting the focal plane until the global NE signal was optimally visualized. The baseplate was affixed using Dental Super Bond Polymer (Hentschel Dental), and a baseplate cap (BCP-2, Inscopix) was placed to protect the lens.

#### Behavioral recordings

After at least one week of recovery, mice were connected to the miniscope system (nVoke2, Inscopix) shortly before behavioral testing and underwent two consecutive days of behavioral assessment. On Day 1, mice performed the established SI task, which consisted of 10 minutes of individual exploration followed by 10 minutes of SI with an unfamiliar juvenile CD1 mouse (4–6 weeks old). On Day 2, the procedure was repeated using an unfamiliar juvenile C57BL/6N mouse (4–6 weeks old) as the social partner. Miniscope recordings were initiated during the final 2 minutes of individual exploration and continued throughout the entire 10-minute SI phase. Behaviors were recorded with a Basler ace2 GigE monochrome camera at 30Hz. Image acquisition and behavior were synchronized using the data acquisition box (DAQ) of the nVoke2 triggered by the Ethovision XT 17 software (Noldus) through a TTL box (Noldus) connected to the USB-IO box from the Ethovision system (Noldus).

#### Data preprocessing

GRAB-NE2h fluorescence was excited with a 470 nm LED and collected in the green channel through a GFP filter set using a head-mounted miniscope coupled to a GRIN lens (Inscopix). Sensor imaging data were processed using the Inscopix Data Processing Software (IDPS, v1.8.0). Videos were first cropped to the usable field of view to remove background regions. Motion correction was then applied to minimize frame-to-frame displacements and prevent artefactual fluorescence fluctuations. Because this experiment used the NE-GRAB sensor providing population-level signals rather than single-cell resolution, no automated cell extraction or component analysis (e.g., PCA/ICA, CNMF-E) was performed. Instead, a single large region of interest (ROI) encompassing the functional field of view was manually defined, and ΔF/F traces were extracted for subsequent analyses.

#### Data analysis of NE signal

We analyzed ΔF/F photometry traces separately for the CD1 and C57 cohorts on a per-recording basis. To account for slow decay/drift in the photometry signal, we next detrended each recording using polynomial regression. For every recording, a fifth-degree polynomial was fit to the ΔF/F time series (as a function of time), and the fitted curve was taken as the baseline trend. The background-corrected trace was obtained by subtracting this trend from the signal. Fits were computed independently per recording (applied column-wise) and the same degree (5) was used for all files, chosen empirically as the optimal degree for these data. The resulting background-corrected traces were used for downstream analyses.

To assess overall difference in NE activity across the whole recording, for each file, the ΔF/F time series was standardized using scipy.stats.zscore with default parameters (axis = 0, ddof = 0), yielding Z-scores computed where μ and σ are the within-recording mean and standard deviation of the trace:

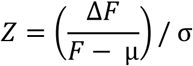

Z-scoring was performed independently for each recording (no pooling across sessions or animals).

Alternatively, to assess differences in NE activity across behavioral events, behavior bouts identified by DeepOF were merged into continuous intervals. An interval was defined from the start frame of the first detected bout to the end frame of the last bout in sequence. Bouts separated by less than 500 ms (corresponding to half of the recording frame rate) were combined into a single interval. Intervals shorter than 500 ms were discarded. In addition, due to the analysis window set to 6 s, intervals occurring less than 6 s apart were excluded to prevent overlap.

Preprocessed ΔF/F signals of NE activity were then aligned to the behavior intervals, using frame-based windows spanning from −3 s before behavior onset to +3 s after onset. These ΔF/F signal intervals were normalized using a robust Z-score (Equation: 1 and 2) with a baseline from the first 2 seconds of the extracted interval. This was performed to mitigate the effects of conditioning or other factors on the signal, and boost the contrast of the behavior component within the signal^73,74^.

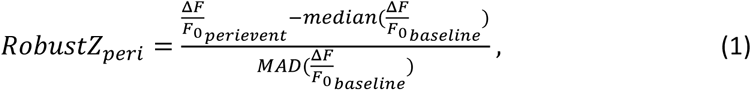

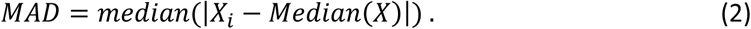

After z-scoring, three preprocessing steps were applied in sequence (enabled by default): (1) outlier suppression, replacing extreme samples with NaN values (apply_outlier_removal=True); (2) gap filling by one-dimensional interpolation to restore continuity across outlier-induced NaNs, allowing propagation from both directions (apply_interpolation=True); and (3) total variation denoising (TVD) to reduce high-frequency noise while preserving step-like changes in the signal (apply_tvd=True). Giving output for each subject×event, a collection of cleaned, z-scored signal segments were retained in long format along with their corresponding event intervals. No across-event or across-subject averaging was done at this stage, as it was included in two separate downstream steps of plotting signal intervals and Area Under the Curve (AUC) signal calculations.

For each subject and event, cleaned z-scored signal segments were used to compute baseline-shifted areas under the signal curve (AUCs). First, a global minimum z-score was identified across all subjects and events and used as a common baseline reference. For each event segment, the first two seconds (corresponding to n = 2 × frame rate samples) after event onset were excluded to minimize onset-related artifacts. The AUC of the remaining segment was then calculated relative to the global minimum baseline. This procedure yielded per-event, per-subject AUC values, which were aligned with subject condition and sex for statistical analysis.

For each behavioral event, average NE signals were plotted separately by condition. Individual signal segments were aligned to event onset and aggregated per condition. The mean signal was computed across subjects, and bootstrap resampling (percentile method, 95% confidence level by default) was used for confidence intervals around the mean. Signals were plotted over a symmetric window centered on event onset (x-axis in seconds).

For each behavioral event, we fit a separate linear mixed-effects model to test the effect of condition on NE activity. The dependent variable was the AUC computed per subject and event (see above). Models included condition as a fixed effect and a random intercept for subject to account for repeated measures within subjects; no random slopes were specified. Prior to modeling, variables were treated as categorical and re-leveled so that WT served as the reference category for condition (comparison: KO vs WT). Parameters were estimated by restricted maximum likelihood (REML) using Python statsmodels’ MixedLM. For each event, we report fixed-effect coefficients, Wald 95% confidence intervals, and p-values. Model fits were saved per event, and results were aggregated across events. If a model failed to converge for a given event, that event was excluded from the summary and the error was logged.

### Statistics

Statistical analyses and graphs were made using Rstudio (with R 4.1.1). All animals were used for statistical analysis unless stated otherwise. Statistical outlier tests were performed using Tukey’s fences method, in which values above quartile (Q) 3 + 1.5 x interquartile range (IQR, calculated as Q3 - Q1) or below Q1 - 1.5 x IQR were considered outliers. This led to the exclusion of 9 animals in the microdialysis experiment, 4 WT and 5 KO mice. Data were tested on the corresponding statistical assumptions, which included the Shapiro-Wilk test for normality and Levene’s test for heteroscedasticity. If assumptions were violated the data were analyzed using non-parametric variants of the test. The two group comparisons were analyzed using the independent samples t-test (T) as a parametric option, Welch’s test (We), if data was normalized but heteroscedastic, or the Wilcoxon test (Wx) as a non-parametric option. The chronic and acute stress exposure data was analyzed using a two-way ANOVA (parametric) or Kruskal-Wallis test (non-parametric) using Chronic and Acute as between subject factors, and further posthoc analysis was performed using the Tukey HSD test (parametric) or the Wilcoxon test (non-parametric). The metabolite measurement data in the microdissection, as well as the microdialysis experiments were analyzed using a two-way ANOVA with the genotype (WT or KO) and the social interaction (NI vs. SI) as between-subject factors. The bar graphs are presented as mean ± standard error of the mean (SEM). Data were considered significant at p<0.05 (*), and further significance was represented as p<0.01 (**), p<0.001 (***), and p<0.0001 (****).

## Acknowledgements

The authors thank Daniela Harbich, Bianca Schmid, Cornelia Flachskamm, and Rainer Stoffel for their technical assistance, Sowmya Narayan and Lea Maria Brix for their experimental support, Marius Fuchs and Prof. Dr. Jochen Klein for the assistance with the microdialysis experiments, Elmira Anderzhanova for the microdialysis sample measurement, Stefanie Unkmeir and Sabrina Bauer of the scientific core unit Genetically Engineered Mouse Models for technical support, and the DeepLabCut development team for creating and maintaining the DeepLabCut software. Furthermore, the authors thank the Max Planck Institute of Biochemistry Mass Spectrometry Core Facility (RRID:SCR_025745) for proteomics sample measurement.

## Author contributions

JB and MVS conceived the study. JB performed the experiments, SC, MR, MP, LD, SN, HY, VK, SM, and MS assisted with the experiments. MDA performed HPLC analysis of microdissection tissue. TB, TE, YM, and NCG performed the proteomics protocol and analysis, JMD provided the genetic mouse line. JB analyzed the data and was assisted by LS, AB, LM, PS, and BMM. JB and MVS wrote the first draft of the manuscript and all authors contributed to the revisions.

## Funding

This study is supported by the “Kids2Health” grant (MVS) of the Federal Ministry of Education and Research (grant number 01GL1743C), the SCHM2360-5-1 grant (MVS) from the German Research Foundation (DFG) and the SAME-NeuroID project (grant number 101079181) of the European Union (MVS).

## Data availability statement

All raw and processed data supporting this study will be made publicly available: miniscope ΔF/F traces and event annotations, DeepOF behavioral labels/metadata at Zenodo; proteomics raw files and search outputs at PRIDE. Processed summary tables are included as Supplementary Data.

## Supplemental materials

**Supplemental figure 1.**
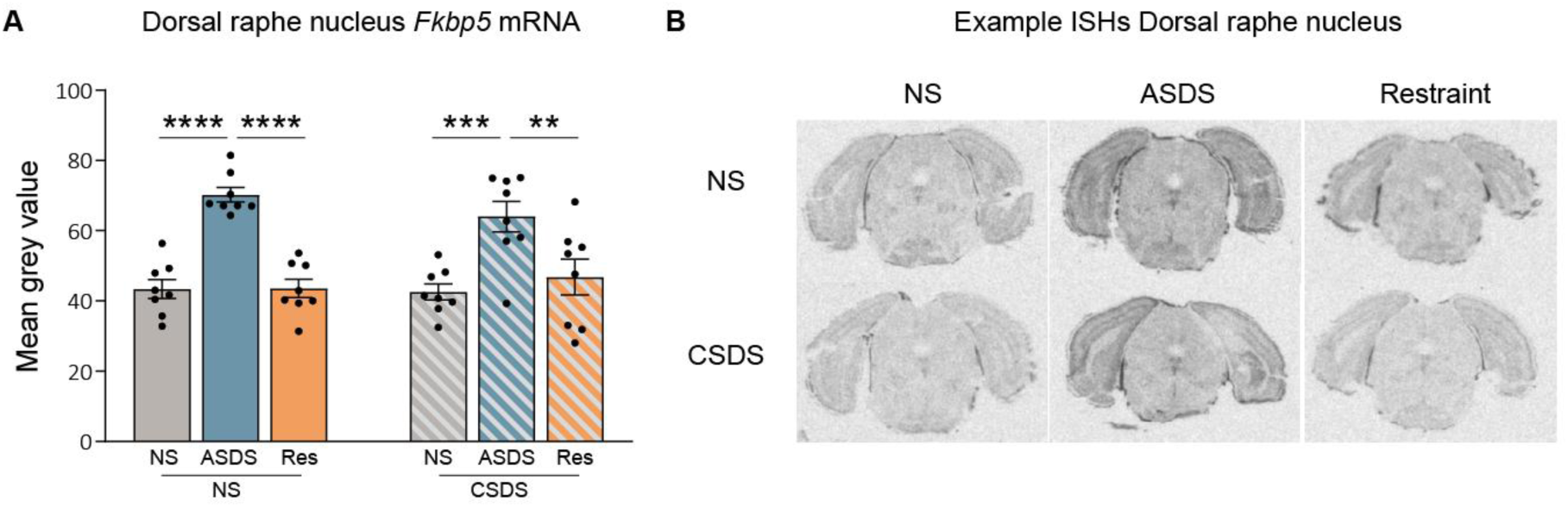
*Fkbp5* expression in the Dorsal raphe nucleus. **A)** *Fkbp5* mRNA in the DRN showed a significant main effect using a two-way ANOVA (Acute x Chronic) of Acute (F(2,42)=31.03, p<0.0001), with no Acute × Chronic interaction (F(2,42)=0.98, p=0.384) and no main effect of Chronic (F(1,42)=0.214, p=0.646). Post-hoc Tukey indicated higher *Fkbp5* in NS-ASDS vs NS-NS (p<0.0001) and vs NS-Res (p<0.0001), and higher CSDS-ASDS vs CSDS-NS (p=0.0008) and vs CSDS-Res (p=0.0103). NS-ASDS did not differ from CSDS-ASDS (p=0.7835). **B)** Example in-situ hybridization scans in the DRN. In panel A; n=8 for all groups.

**Supplemental figure 2.**
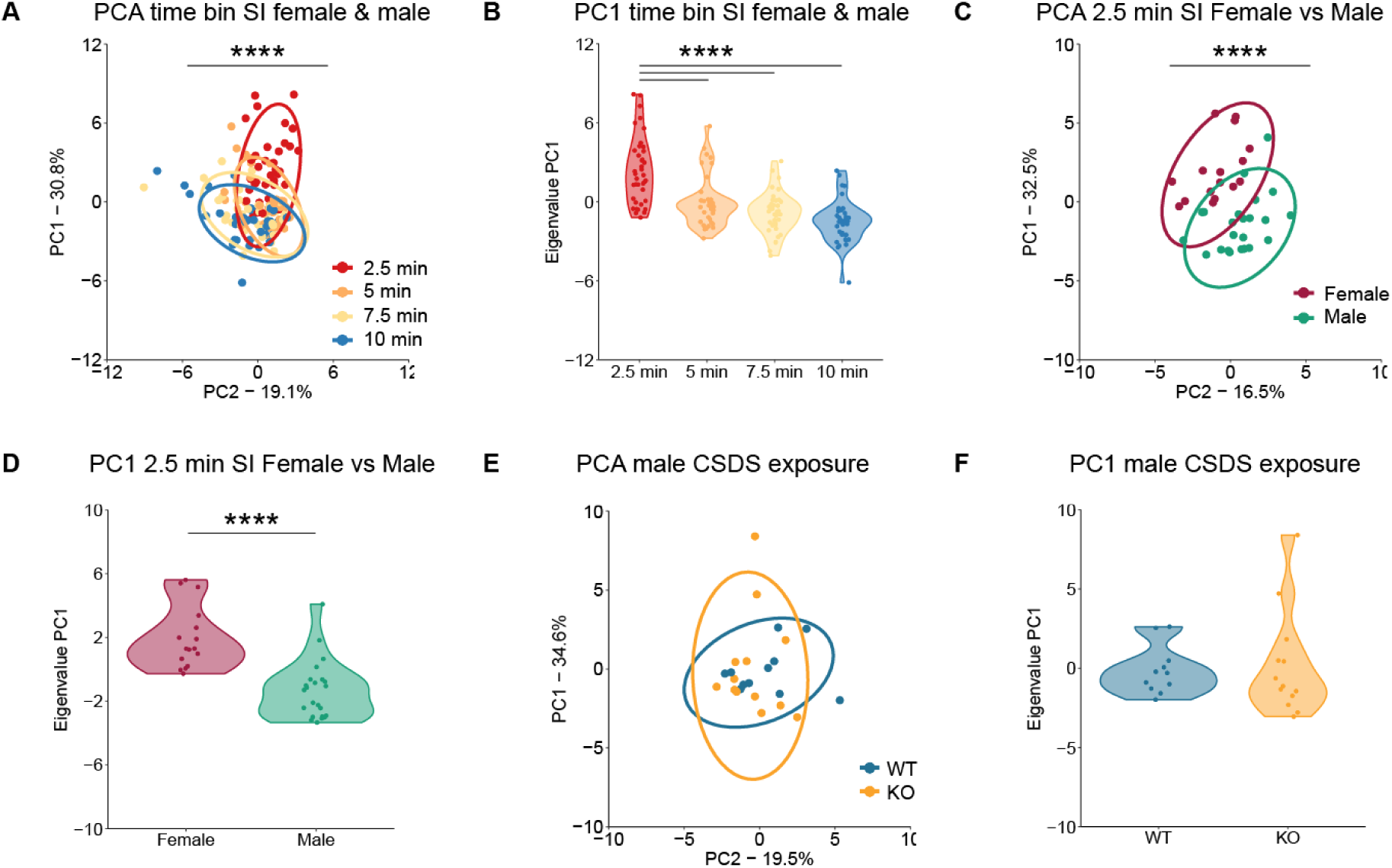
PCA analysis for the female and male social interaction task. **A)** The PCA time bin analysis comparing 4 bins revealed that the first 2.5 min time bin is significantly different from the other time bins. (Kruskal-Wallis test: H(3)=56.00, p=4.21^e-12^. **B)** The PC1eigenvalues of the SI time bin PCA. Post-hoc Wilcoxon: 2.5min vs. 5 min (W=1191, p=8.72^e-6^), 2.5 min vs. 7.5 min (W=1321, p=1.91^e-9^), 2.5 min vs. 10 min (W=1415, p=2.33e-13). **C-D)** The PCA on the 2.5 min SI data revealed a significant difference between sexes (Two-tailed Wilcoxon test: W=339, p=1.17^e-6^). **E-F)** The PCA on the 2.5min SI data for males showed no differences between WT and KO after CSDS exposure (Two-tailed Wilcoxon test: W=82, p=0.57). In panels A-B; n=39 across all four-time bins. In panels C-D; n=16 for female and n=23 for male. In panels E-F; n=11 for WT and n=13 for KO.

**Supplemental figure 3.**
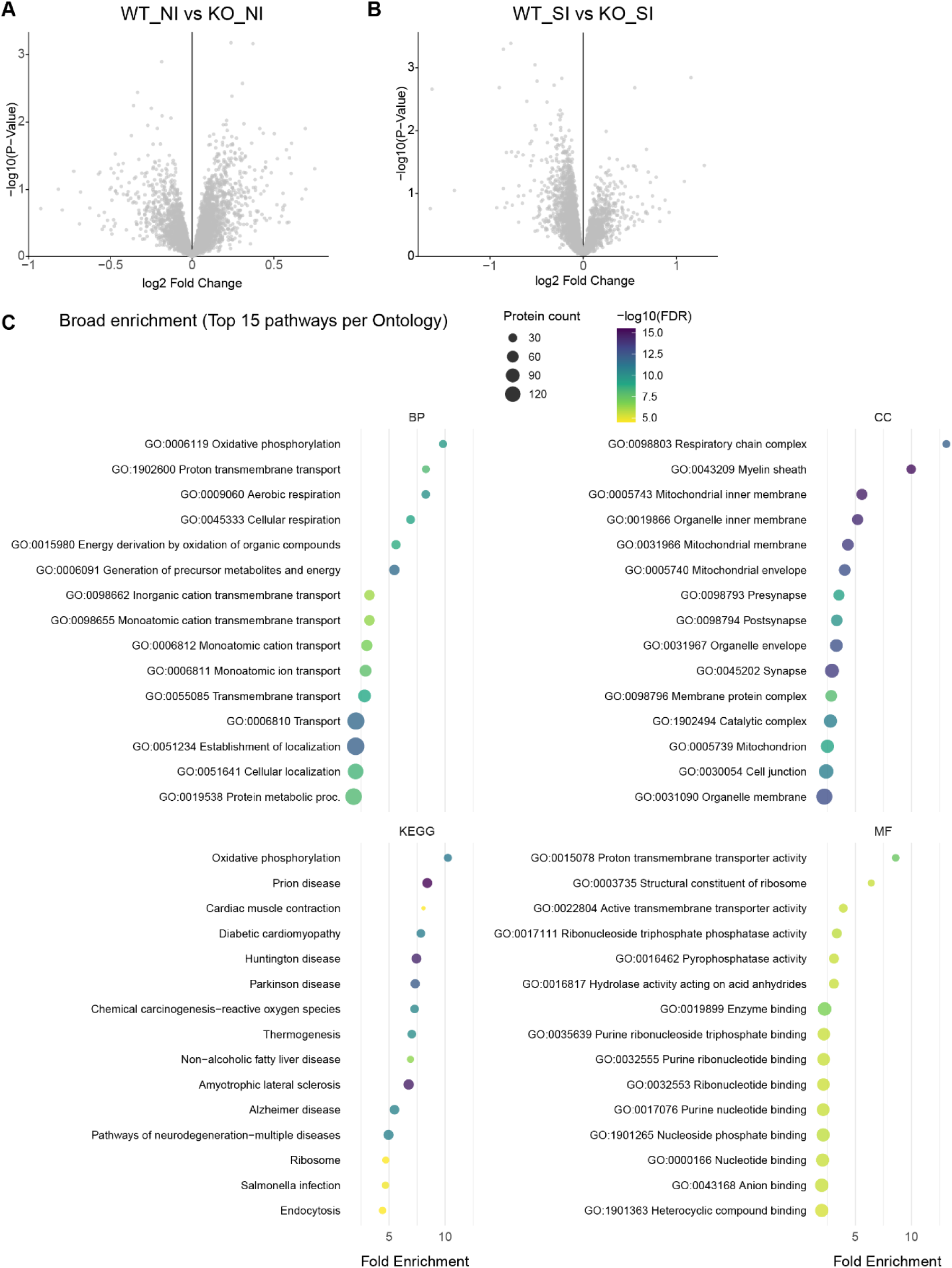
Proteomic analysis of social interaction in WT and KO mice. **A)** Volcano plot showing the down- and upregulated proteins in WT_NI vs KO_NI (Adj p-value<0.05: down: 0; Up: 0). **B)** Volcano plot showing the down- and upregulated proteins in WT_SI vs KO_SI (Adj p-value<0.05: down: 0; Up: 0). **C)** The proteomics GO TERM and KEGG analysis between WT and KO after NI and SI showing the top 15 molecular pathways. Sample size: panels A-C: WT_NI: n=10, WI_SI: n=10, KO_NI: n=11, and KO_SI n=10.

**Supplemental figure 4.**
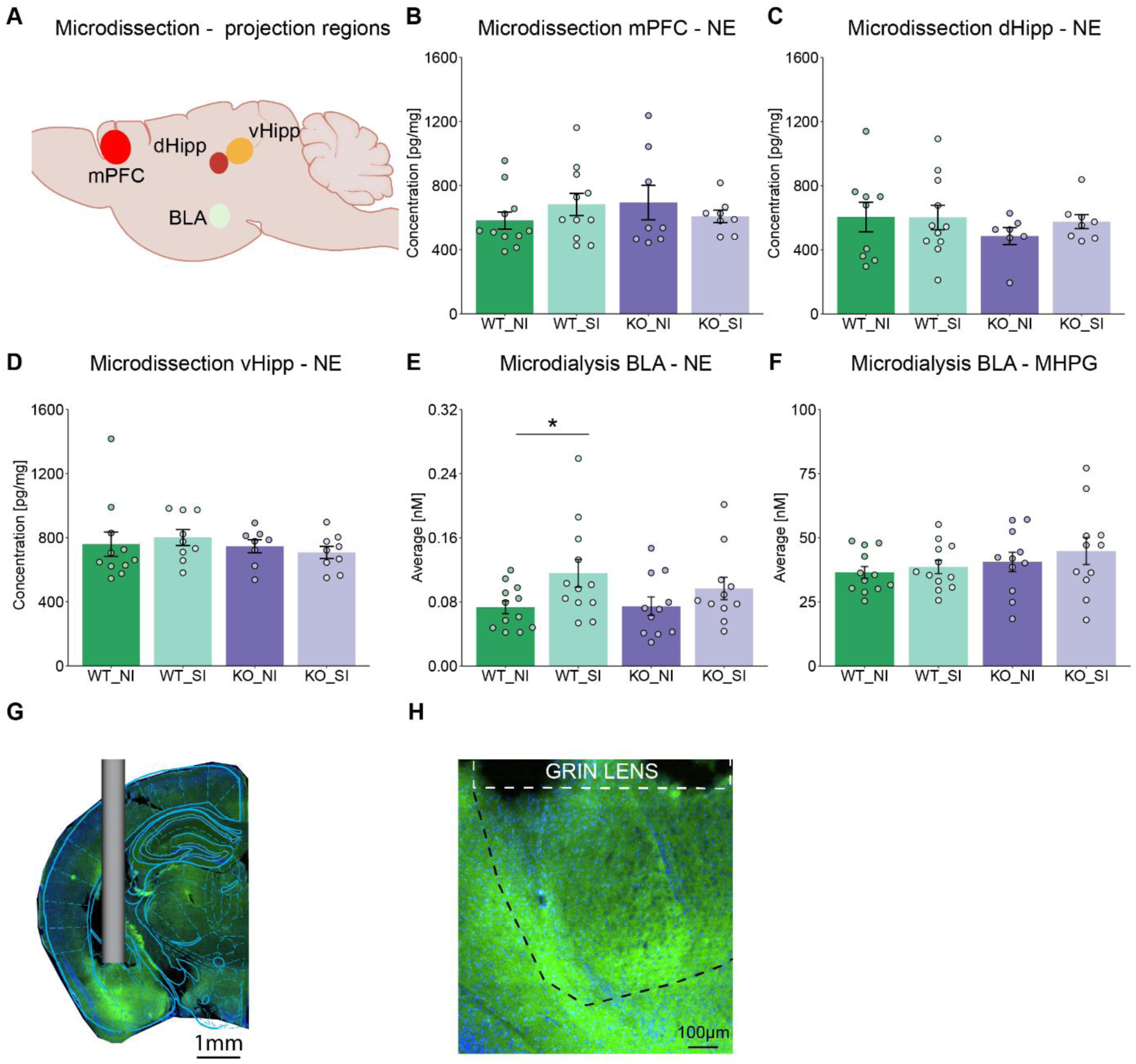
Noradrenergic signaling between different project regions. **A**) Schematic overview of analyzed project regions of the LC with microdissection **B)** Microdissection measuring total NE levels in the mPFC. No main effect differences were observed using the two-way ANOVA: social interaction: F(1,34)=0.1, p=0.75, genotype: F(1,34)=0.07, p=0.79, social interaction*genotype: F(1,34)=1.73, p=0.20. **C)** Microdissection measuring total NE levels in the dHipp. No main effect differences were observed using the two-way ANOVA: social interaction: F(1,31)=0.27, p=0.61, genotype: F(1,31)=0.86, p=0.36, social interaction*genotype: F(1,31)=0.38, p=0.54. **D)** Microdissection measuring total NE levels in the vHipp. No main effect differences were observed using the two-way ANOVA: social interaction: F(1,33)=0.0001, p=0.99, genotype: F(1,33)=0.81, p=0.37, social interaction*genotype: F(1,33)=0.50, p=0.49. **E)** Microdialysis measuring NE release in the BLA. Posthoc Tukey revealed that WT social interaction (WT_SI) has significantly increased NE concentrations in the BLA compared to WT no interaction (WT_NI), F(1,22)=4.99, p=0.036, which was not altered in the *Fbkp5*^Nat^ background, comparing KO_NI with KO_SI, p=0.23. Two-way ANOVA: social interaction; F(1,42)=6.22, p=0.017, genotype; F(1,42)=0.46, p=0.50, social interaction*genotype; (F(1,42)=0.59, p=0.45. **F)** Microdialysis measuring MHPG release in the BLA. No main effects were observed using the two-way ANOVA: social interaction; F(1,42)=0.75, p=0.39, genotype; F(1,42)=2.03, p=0.16, social interaction*genotype; F(1,42)=0.085, p=0.77. In panel B; n=11 for WT_NI, n=11 for WT_SI, n=8 for KO_NI, and n=8 for KO_SI. In panel C; n=9 for WT_NI, n=11 for WT_SI, n=7 for KO_NI, and n=8 for KO_SI. In panel D; n=11 for WT_NI, n=9 for WT_SI, n=8 for KO_NI, and n=9 for KO_SI. In panels E-F; n=12 for WT and n=11 for KO. **G)** Overview image with schematic overlay and included location of the guide cannula (microdialysis) or GRIN lens (miniscope). **H)** Zoom in image of the BLA with the injection site for miniscope NE signaling.

**Supplemental figure 5.**
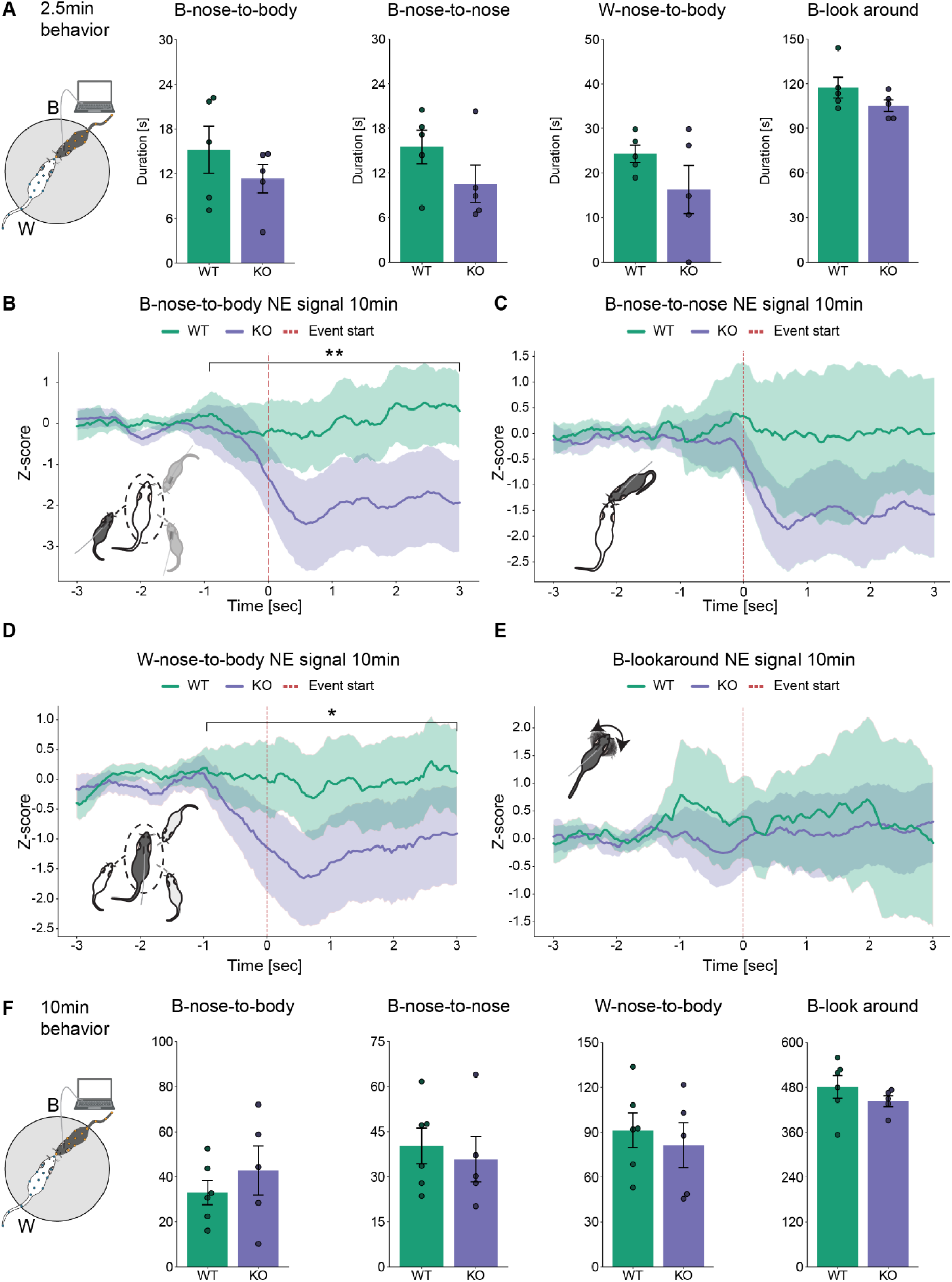
*Fkbp5*^Nat^ deletion remains disrupted during 10min social interaction of BLA NE dynamics. **A)** Schematic overview of the test animal (B) connected to in-vivo NE imaging during social interaction with an outbred CD1 conspecific (W). No significant differences were observed between WT and KO during the first 2.5min total duration of the behaviors, B-nose-to-body (T(8) = −1.05, p = 0.324), B-nose-to-nose (T(8) = −1.46, p = 0.181), W-nose-to-body (T(8) = −1.40, p = 0.200), B-lookaround (T(8) = −1.52, p = 0.167). **B-E)** Event-aligned NE signal traces (±3 s) around specific DeepOF identified social behaviors; **B**) The 10 min NE signal during B-nose-to-body in KO mice showed a significant reduction compared to WT (Mixed linear model β₍KO₎ = −195.61 ± 70.01, z = −2.79, p = 0.005). **C)** The 10 min NE signal during B–nose-to-nose in KO mice did not differ from WT (Mixed linear model β₍KO₎ = −138.47 ± 105.63, z = −1.31, p = 0.190). **D)** The 10 min NE signal during W-nose-to-body in KO mice showed a significant reduction compared to WT (Mixed linear model β₍KO₎ = −124.98 ± 54.57, z = −2.29, p = 0.022). **E)** The 10 min NE signal during B-lookaround in KO mice did not differ from WT (Mixed linear model β₍KO₎ = −35.85 ± 71.06, z = −0.51, p = 0.614). **F)** No significant differences were observed between WT and KO during the total 10min duration of the behaviors, B-nose-to-body (T(9) = 0.84, p = 0.421), B-nose-to-nose (T(9) = −0.46, p = 0.654), W-nose-to-body (T(9) = −0.54, p = 0.605), B-lookaround (T(9) = −1.05, p = 0.321). Sample size: panels A-F: WT: n=5 and KO n=5.

**Supplemental figure 6.**
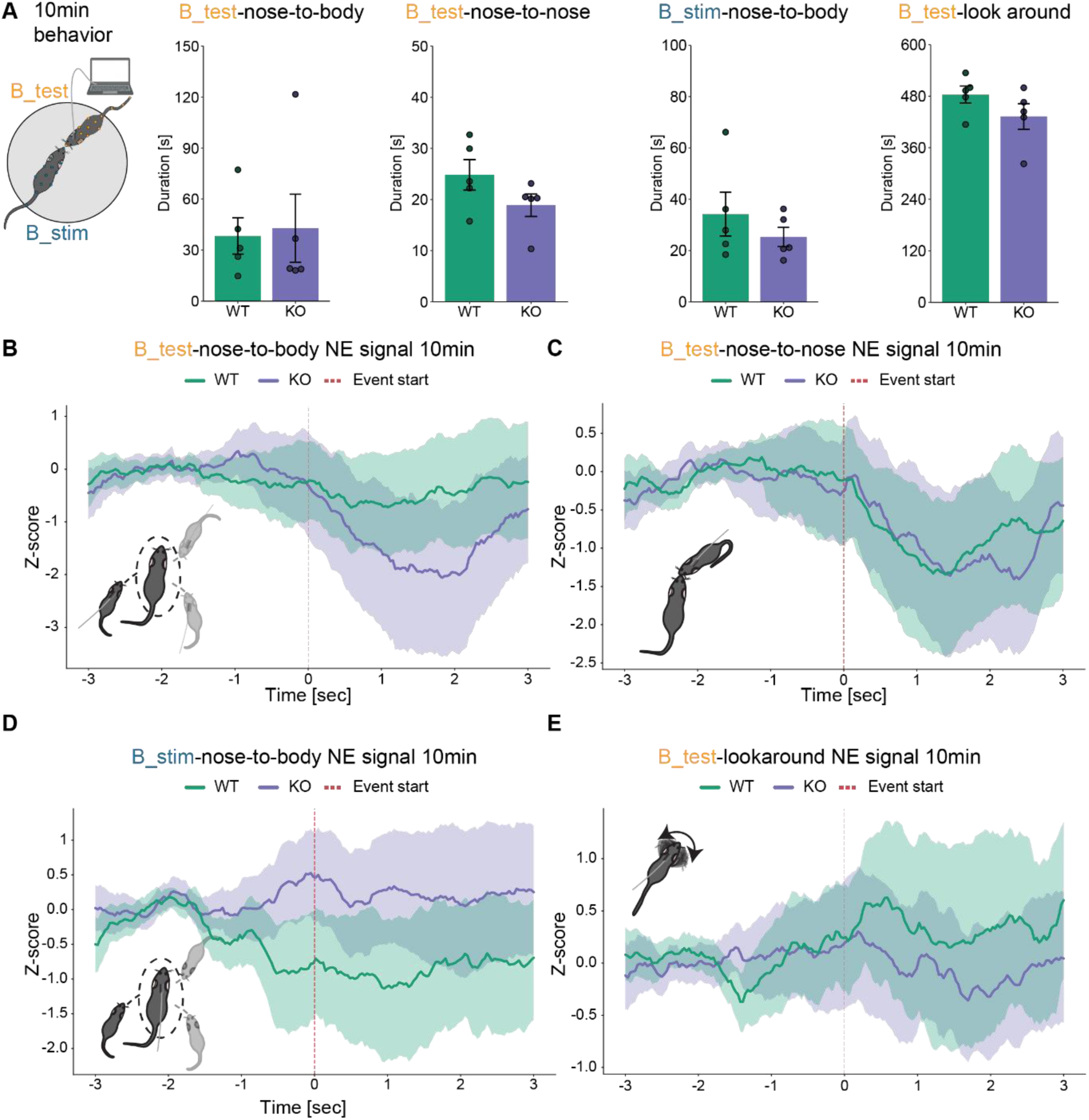
*Fkbp5*^Nat^ deletion remains not disrupted during 10min same strain C57/Bl6N social interaction of BLA NE dynamics. **A)** Schematic overview of the test animal (B_test) connected to in-vivo NE imaging during social interaction with an outbred CD1 conspecific (B_stim). No significant differences were observed between WT and KO during the total 10min duration of the behaviors, B-nose-to-body (T(8) = 0.20, p = 0.846), B-nose-to-nose (T(8) = −1.60, p = 0.149), W-nose-to-body (T(8) = −0.96, p = 0.364), B-lookaround (T(8) = −1.43, p = 0.190). **B-E)** Event-aligned NE signal traces (±3 s) around specific DeepOF identified social behaviors; **B)** The 10-min NE signal during B_test-nose-to-body in KO mice did not differ from WT (Mixed linear model β₍KO₎ = −78.94 ± 87.05, z = −0.91, p = 0.365). **C)** The 10-min NE signal during B_test-nose-to-nose in KO mice did not differ from WT (Mixed linear model β₍KO₎ = −8.41 ± 66.50, z = −0.13, p = 0.899). **D)** The 10-min NE signal during B_stim–nose-to-body in KO mice did not differ from WT (Mixed linear model β₍KO₎ = 124.45 ± 64.92, z = 1.92, p = 0.055). **E)** The 10-min NE signal during B_test-lookaround in KO mice did not differ from WT (Mixed linear model β₍KO₎ = −36.28 ± 50.91, z = −0.71, p = 0.476).

## Notes

### Competing Interest Statement

The authors have declared no competing interest.

